# Modeling Somatic Second-Hit Mutations in Novel Mouse Models of Hereditary Hemorrhagic Telangiectasia

**DOI:** 10.64898/2026.02.15.706014

**Authors:** Adella P. Bartoletti, Shreya Bavishi, K C Rajan, Stryder M. Meadows

**Author notes:** **Corresponding Author**: Stryder M. Meadows, **Correspondence Email:**.

## Abstract

Hereditary Hemorrhagic Telangiectasia (HHT) is a genetic vascular disorder characterized by distinct vascular malformations, including deep organ arteriovenous malformations (AVMs) and mucocutaneous telangiectasias. People with HHT inherit monoallelic pathogenic variants in members of the TGFβ signaling cascade (*ACVRL1*, *ENG* and *SMAD4*), resulting in a loss of gene function and dysangiogenesis. While these heterozygous inactivating mutations are present in all cells, malformations develop locally, indicating a focal trigger of onset. Indeed, recent human sequencing studies revealed that second-hit somatic mutations, resulting in complete bi-allelic loss of gene function, are linked to lesion formation in the three major types of HHT (HHT1, HHT2, JP/HHT). To model the loss of heterozygosity (LOH) associated with HHT patients, we generated new Eng and Smad4 HHT mouse models whereby endothelial cell-specific, somatic LOH mutations are induced within a heterozygous loss of function background (*HHT-*iEC-LOH). The *HHT-*iEC-LOH models recapitulate the mosaic makeup of patient malformations and indicate that multiple, distinct secondary somatic mutations can contribute to AVM onset. Utilizing immunofluorescent staining, blue latex vasculature casting, weighted tracer perfusions, and lineage tracing studies, *HHT-*iEC-LOH models were phenotypically assessed and compared to traditional inducible endothelial cell knockout (*HHT-iECKO*) HHT models. Overall, *HHT-*iEC-LOH mice exhibit increased malformation frequency and vascular phenotypes that are comparable or exceed the severity of iECKO models. Significantly, *HHT-*iEC-LOH mice can be induced early in development and live into adulthood, displaying persistent cerebrovascular phenotypes. The heightened patient representation offered by these newly developed models enables the study of long-term disease progression and testing of therapeutic interventions.

## Introduction

Hereditary Hemorrhagic Telangiectasia (HHT) is a rare, heritable genetic disorder in which patients develop vascular malformations, excessive epistaxis, and anemia[1, 2]. While HHT has long been categorized as a rare disease (1 in 5,000 people), recent publications suggest its prevalence may be twice as high, or more, than originally estimated[3, 4]. Vascular malformations in HHT consist of surface-level entangled vessels (telangiectasias) and enlarged, fragile connections in deep organs that bypass the normal capillary network (arteriovenous malformations, AVMs). These enlarged, weak vessels can rupture, causing severe bleeding, including in major organs or from mucosal surfaces (i.e., nosebleeds) [1, 2, 5, 6]. Generalized bleeding greatly affects HHT patient quality of life, with many reporting that the fear of bleeding or sudden onsets of bleeding have prevented daily tasks[7]. Vascular defects from HHT can also result in other dangerous complications, such as stroke and even death[1–4]. Though our knowledge of the various mechanisms behind the pathogenesis of HHT is expanding, therapeutic interventions have been unable to fully treat malformations and have been more beneficial in respect to daily nosebleeds[8–13]. Thus, further research is needed to understand the interplay between signaling cascades and cellular communication leading to malformation development and vessel integrity defects that cause consistent bleeding problems and life-threatening consequences.

People with HHT are born with inherited pathogenic variants in one of three members of the TGF-β signaling family – *Endoglin* (*ENG, HHT1*), *Activin receptor-like kinase-1* (*ALK1, HHT2*) or *Smad-related protein 4* (*SMAD4, JP-HHT*) – resulting in mono-allelic loss of function (haploinsufficiency) of the affected gene[1, 2, 5, 6, 14–20]. While this heterozygous inactivation is systemic, vascular malformations form focally. AVMs often form in visceral organs (lung, liver, and brain), and telangiectasias often form on areas of the skin that experience friction[1, 14]. In all HHT variations, the localized formation of lesions suggests a focal trigger of onset in the endothelium. Based on Knudson’s two-hit theory, it has been hypothesized that a second hit, such as environmental triggers or random somatic mutations, leads to loss of function of the remaining functional allele and subsequent loss of heterozygosity (LOH), which drives localized malformation[21]. Supporting this notion, patient sequencing studies have recently shown that some telangiectasias and AVMs across all three HHT types are mosaic, consisting of cells with both mono-allelic and bi-allelic loss of function (LOH)[22–26]. These studies demonstrated that secondary mutations occurred at different locations within the “functional” gene, on the allele without the germline mutation, were distinct in malformations from the same individual, and were absent in constitutive DNA collected from non-lesion areas. These findings strongly indicate that local, random somatic mutations drive the mosaic formation of bi-allelic loss of function lesions in HHT patients.

Currently, no available animal models accurately recapitulate the mosaic pattern of biallelic loss of function seen in the heterozygous HHT patients’ genetic background. Available mouse models for HHT are generally heterozygous, with one completely wild type allele and one inactivated allele, or inducible homozygous models, with two active alleles that can be knocked out in the endothelial cells (EC) after birth using the Cre-loxP system[27–38]. Unfortunately, heterozygous mouse models of HHT do not consistently form distinctive lesions[27, 32]. In contrast, homozygous inducible mouse models form robust AVMs during postnatal developmental angiogenesis or require additional stimulation with wounding[39, 40] or VEGF injections at adulthood[41, 42] (embryonic induction leads to *in utero* lethality). These requirements offer support for the influence of second hits on local AVM formation but may also be indicative of potential weaknesses with these models, as they do not take into account the heterozygous genetic background. Furthermore, early postnatal induction of genetic inactivation in inducible homozygous HHT models leads to lethality within 2-10 days depending on the model[30, 39, 40, 43], while gene inactivation in adults generally does not result in AVM formation[44]. These outcomes severely inhibit the ability to test the effects of therapeutics long-term while monitoring whole body effects, which is more applicable to patients. To circumvent early lethality, researchers have induced local gene deletion during the neonatal period using either tissue-targeted tamoxifen injections or brain-specific Cre drivers to promote vascular lesion development[45, 46]. Nonetheless, these models still do not recapitulate the heterozygous genetic nature of the disease and are limited to a specific tissue or region of the body. Therefore, there is a critical need to create and utilize HHT animal models in which LOH can be induced during developmental angiogenesis, similar to HHT patients, and its effects as a second hit explored longitudinally. This more accurate depiction of the genetic makeup of HHT is necessary to better understand the EC-EC communication and molecular and cellular pathways involved in HHT vascular lesion pathogenesis.

The studies presented herein aimed to address limitations in HHT research by generating new HHT mouse models that incorporate the ability to induce somatic mutations specifically in ECs and in the context of heterozygous HHT backgrounds (LOH). We established *Eng*- and *Smad4*-iEC-LOH HHT mouse models that develop mosaic, biallelic loss of function lesions, similar to HHT patients. Utilizing various phenotypic characterization methods, we detailed vascular lesion prevalence, morphology, and blood vessel integrity in comparison to the established EC-inducible homozygous *Eng* and *Smad4* mice. This includes a description of AVM formation and vessel enlargement in the retina and brains during neonatal angiogenesis. Further, we show that the LOH models can survive into early adulthood and present persistent cerebrovascular defects, thereby establishing a longitudinal model to study HHT phenotypes. Together, these novel LOH HHT models may provide critical insights into the cellular mosaicism and development of AVMs and establish a long-term platform to assess therapeutic avenues for treating HHT.

## Materials and Methods

### Mice and Treatments

All animal experiments were conducted in accordance with Tulane University’s Institutional Animal Care and Use Committee. To establish both *HHT*-iEC-LOH (HHT = *Smad4* or *Eng*) models, *HHT^f/f^* (*Smad4*[47] *–* Jax Lab Strain 017462; *Eng*[28]) mice were crossed with epiblast-specific *Sox2Cre* mice[48] (Jax Lab Strain 008454) to generate mono-allelic knockout of *HHT* (*HHT^f/wt^;Sox2Cre*, which is equivalent to *HHT^-/wt^;Sox2Cre*). These mice were then backcrossed with *HHT^f/f^*mice to generate *HHT^f/-^* mice without *Sox2Cre* (Sup Fig 1a,b). *HHT^f/-^* mice were further crossed with *Cdh5*-*Cre^ERT2^*mice[49] to generate endothelial cell (EC) specific, tamoxifen inducible heterozygous knockout mice: *HHT^f/-^;Cdh5*-*Cre^ERT2^*(otherwise referred to as ***HHT*-iEC-LOH**, Sup Fig. 1c,d). To track Cre expression and evaluate the cellular makeup of an AVM, two reporter strains – *mTmG*^+/+^[50] (Jax Lab Strain 007676) and *confetti*^+/+^[51] (Jax Lab Strain 017492) – were incorporated into the *HHT-iEC-LOH* mouse lines, generating *HHT;mTmG^+/-^ ;iEC-LOH* and *HHT;confetti^+/-^;iEC-LOH* mice (Sup Fig. 1e,f) for analysis.

For characterization experiments, *HHT^f/-^;Cdh5-Cre^ERT2^*mice were mated with *HHT^f/f^* mice to generate four sample groups – *HHT^f/-^* and/or *HHT^f/f^*, with and/or without *Cdh5-Cre^ERT2^* (Sup Fig 1d). For AVM makeup studies, *HHT^f/-^;Cdh5-Cre^ERT2^*mice were mated with *HHT^f/f^;mTmG^+/+^* mice to generate four sample groups: *HHT^f/-^;mTmG^+/wt^* and *HHT^f/f^;mTmG^+/wt^*, with or without *Cdh5-Cre^ERT2^* (Sup Fig 1e). To further the cellular makeup studies, *HHT^f/-^;Cdh5-Cre^ERT2^* mice were mated with *HHT^f/f^;confetti^+/+^* mice to generate four sample groups: *HHT^f/-^; confetti^+/wt^* and *HHT^f/f^;confetti^+/wt^*, with or without *Cdh5-Cre^ERT2^*(Sup Fig 1f).

To induce recombination of the target sequence (detailed strategy depicted in Sup Fig. 2a and d), tamoxifen (Sigma) was dissolved in 100% ethanol and sunflower oil (1:10) and orally fed with a pipette tip to neonatal pups at designed postnatal (P) days based on previous studies[52]. For comparison of the new *Smad4*-iEC-LOH model to the previous iECKO model and determining the cellular makeup of an AVM, 5 µg of tamoxifen was delivered orally at P1 to induce partial EC deletion of *Smad4* (Sup Fig 2c). For comparison of the new *Eng-*iEC-LOH model to the previous iECKO model and determining the cellular makeup of an AVM, 2.5 µg of tamoxifen was delivered orally at P1, to induce partial EC deletion of *Eng* (Sup Fig 2f). All mice were collected at P7 or at 10 weeks of age (P70) (Sup Fig 2c and f). The sex of the mice was not distinguished in most of these studies; prevalence of HHT affects males and females equally[53].

### Genotyping

Tissue for genotyping was collected from mice by toe clipping at P1, and the representative genotyping strategies and gels are shown in Sup Fig. 2a, b, d, e. The knockout gene (-) must be genotyped prior to tamoxifen treatment, as the endothelial loss of *Smad4* or *Endoglin* following tamoxifen induction of Cre can be enough to show up as a noticeable gel band. Genotyping of the KO and *Cdh5* alleles included a primer mix for *vascular endothelial growth factor* (*VEGF*) as an internal control to verify individual genotyping success.

Mice were genotyped using Quanta AccuStart II master mix with the following primers:

Smad4 Flox Forward: TAA GAG CCA CAG GGT CAA GC;

Smad4 Flox Reverse: TTC CAG GAA AAA CAG GGC TA;

Smad4 KO Forward: CAG AGT GGG TCT TTC TAC CTT;

Smad4 KO Reverse: GAC CCA AAC GTC ACC TTC AC;

Eng Flox Forward: GTG GTT GCC ATT CAA GTG TG;

Eng Flox Reverse: GGT CAG CCA GTC TAG CCA AG;

Eng KO Forward: GGT CAG CCA GTC TAG CCA AG;

Eng KO Reverse: CCA CGC CTT TGT CCT TGC;

Cdh5-Cre^ERT2^ Forward: TCC TGA TGG TGC CTA TCC TC;

Cdh5-Cre^ERT2^ Reverse: CCT GTT TTG CAC GTT CAC CG;

mTmG Wild Type Forward: AGG GAG CTG CAG TGG AGT AG;

mTmG Mutant Forward: TAG AGC TTG CGG AAC CCT TC;

mTmG Reverse: CTT TAA GCC TGC CCA GAA GA;

confetti Wild Type Forward: AAA GTC GCT CTG AGT TGT TAT;

confetti Mutant Forward: GAA TTA ATT CCG GTA TAA CTT CG;

confetti Reverse: CCA GAT GAC TAC CTA TCC TC;

SRY Forward: TTG TCT AGA GAG CAT GGA GGG CCA TGT CAA;

SRY Reverse: CCA CTC CTC TGT GAC ACT TTA GCC CTC CGA;

VEGF Forward: CCT GGC CCT CAA GTA CAC CTT;

VEGF Reverse: TCC GTA CGA CGC ATT TCT AG;

Cre Forward: GAT CGC TGC CAG GAT ATA CG;

Cre Reverse: CAT CGC CAT CTT CCA GCA G;

Sox2-Cre Wild Type Forward: CTT GTG TAG AGT GAT GGC TTG A;

Sox2-Cre Mutant Forward: CCA GTG CAG TGA AGC AAA TC;

Sox2-Cre Reverse: TAG TGC CCC ATT TTT GAA GG.

Genotyping polymerase chain reactions were run on a BioRad C1000 Touch or T100 touch thermal cycler utilizing previously described protocols. The resulting DNA was run on 2% agarose gel with 3.5% ethidium bromide and imaged using a BioRad GelDoc XR+.

### Litter Analysis

Neonate mice were genotyped as above, with the addition of SRY primers to determine the number of males and females in each litter. At P7, mice body and heart weights were measured using a Sartorius Quintix612-1S after euthanasia. Heart size was determined as a percentage of total body weight: (heart weight + body weight) x 100.

### Fluorescent-Weighted Tracer Perfusion

After anesthetizing neonate and mice using isoflurane (VetOne Fluriso), abdominal and thoracic cavities were opened to expose the hearts. At P7, combined fluorescent tracers: 10kDa dextran (FITC-conjugated) at 20mg/mL (Sigma-Aldrich) and 70kDa dextran (Texas-red conjugated) at 10mg/mL (Thermoscientific) were perfused into the left ventricle using a 27-gauge needle with a 1mL syringe: 800 μL total (400 μL of each tracer). Tracers were allowed to circulate directly through the arterial system for approximately 10-15 minutes, then the right atria were cut (Sup Fig 6a). Following tracer perfusion, 5 mL of phosphate buffered saline (PBS) was perfused using a 27-gauge needle with a 10 mL syringe to clear any tracer leftover in the vessels (Sup Fig 6b), allowing only tracers that leaked out of the vessels to remain (Sup Fig 6c). A syringe setup with a 2-way syringe connector was used to avoid removing and reinserting the needle between tracer and PBS perfusions. Mice were then decapitated, the skin on the head was pulled away to expose the skull for removal, and brains were placed in PBS and immediately imaged using a Leica M205 FA stereomicroscope. Analysis of tracer presence in the brain was completed by denoting presence (100) or absence (0). Eyes from these mice were also collected and dissected (as explained below) to stain for red blood cell presence (due to leakage) following PBS clearance of the vascular beds. Fluorescent tracers were not observed in eyes following collection and processing, which was expected because the tracers used for this study were not PFA fixable and were likely washed away with retina preparation. However, *Ter119* was marked with a far red secondary to avoid any potential background from the tracers.

### Retinal Dissection and Preparation

For *Ter119* staining of red blood cells, mice were perfused as stated above. For Isolectin IB4 and *ERG* staining, mice were not perfused. At P7, mice were euthanized through decapitation, and eyes were collected and fixed in 4% paraformaldehyde (PFA) for 1 hour at 4°C. Following fixation, eyes were washed with PBS and dissected. To isolate the retina, the outer sclera was first removed by piercing the cornea and pulling away the outer white layer. Next, the iris and lens were removed followed by plucking the hyaloid vessels from the center of the retina[54].

### Retinal Immunofluorescent Staining

Following retina dissection, retinas were immediately stained for imaging. First, retinas were washed 3 times in PBS for 5 minutes on a nutator at room temperature. Next, retinas were permeabilized with 1% Triton-X dissolved in PBS for 30 minutes at room temperature. Following permeabilization, epitopes were blocked using CAS-Block (Invitrogen) for 30 minutes at room temperature. The retinas were then placed in primary antibodies diluted in 1% Triton-X dissolved in PBS and incubated overnight at 4°C on a nutator. For conjugated primary antibodies, the retinas were then washed 3 times in PBS for 5 minutes on a nutator at room temperature and then flat mounted in Prolong Diamond Antifade Mountant (Invitrogen). For primary antibodies that were not conjugated, retinas were washed 3 times in PBS for 5 minutes on a nutator at room temperature. Secondary antibodies, diluted in 1% Triton-X dissolved in PBS, were then applied for 4 hours on a nutator at room temperature. Following incubation, retinas were washed 3 additional times (5 minutes each) in PBS and flat mounted in Prolong Diamond Antifade Mountant (Invitrogen). All retinas were immediately imaged using a Nikon Eclipse Ti2 confocal microscope. Images were viewed and LUTs were adjusted equally among samples with Nikon Elements. Analysis was completed using Fiji ImageJ Software.

The following primary antibodies were used:

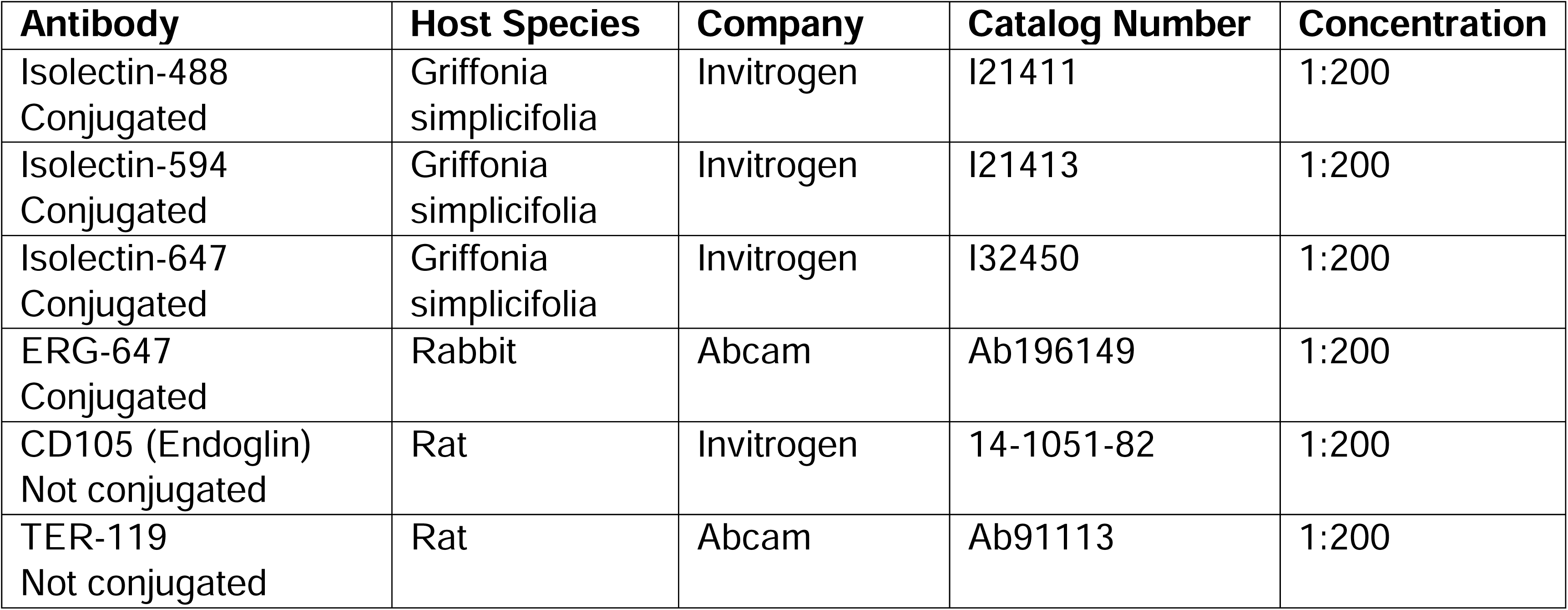

The following secondary antibodies were used at a 1:500 concentration:

| **Fluorophore** | **Host Species** | **Target Species** | **Company** | **Catalog Number** |
| --- | --- | --- | --- | --- |
| 633 | Goat | Rat | Invitrogen | A21094 |

### Brain Collection for Hemorrhage Analysis

P70 experimental mice were anesthetized using isoflurane and then euthanized by decapitation. After removing the skull, brains were fixed in 4% PFA overnight at 4°C, followed by washing with PBS and storage in PBS with 0.01% sodium azide (NaN_3_) at 4°C. Brains were positioned in PBS and imaged using a Leica M205 FA stereomicroscope.

### Blue Latex Dye Perfusion

After anesthetizing mice using isoflurane, abdominal and thoracic cavities were opened to expose the hearts. Holden’s blue latex dye (Reynolds Advanced Materials) was perfused into the left ventricle using a 27-gauge needle with a 1mL syringe. For pups at P7, 1mL of latex was perfused; for adults, 5 mL of latex was perfused by refilling the 1mL syringe but keeping the needle inserted. Perfusion into the left ventricle allowed the dye to circulate directly through the arterial system. After removing the skull, latex-perfused brains were fixed in 4% PFA overnight at 4°C, followed by washing with PBS and storage in PBS with 0.01% sodium azide (NaN_3_) at 4°C. Brains were positioned in PBS and imaged using a Leica M205 FA stereomicroscope. Analysis was completed using Fiji ImageJ Software.

### Statistical Analysis

Sample size (N) values equate to biological replicates (number of mice included in the experiment). All statistical analyses were performed using Graph Pad Prism, Version 10. Bar graphs with mean ± SEM, shown as error bars, were presented for quantified data. Prior to comparison of sample groups, data sets were tested for normal distribution using the “Normality and Lognormality Tests”. Specifically, the distribution was deemed parametric or non-parametric using the Kolmogorov-Smirnov test, as some sample sizes were less than 10. If the distribution was normal, an ordinary one-way ANOVA was used to test significance between more than two groups. If any data set was nonparametric, the Kruskal-Wallis test (nonparametric ANOVA) was used to test significance between more than two groups. As every comparison is independent, no compensation for multiple comparisons was made. A P-value ≤0.05 was considered significant.

## Results

### Generating novel mouse models of HHT that more closely resemble biallelic loss in a heterozygous genetic background

People affected with HHT inherit monoallelic loss of function mutations, making them heterozygous for expression of the HHT related genes *ENG*, *ACVRL1/ALK1* or *SMAD4*[18–20] (Fig 1a). Recent sequencing studies demonstrate that second hits, such as somatic inactivating mutations in the corresponding wildtype allele, can result in localized, biallelic loss of function due to LOH and contribute toward malformation development[22–26]. However, current HHT inducible <u>e</u>ndothelial <u>c</u>ell-specific <u>k</u>nock <u>o</u>ut (*HHT-iECKO*) models do not replicate this genetic progression. Mice are born with intact gene expression, and homozygous deletion is induced postnatally to promote vascular phenotypes, such as AVM formation and vessel enlargements (Fig 1a). To better represent the HHT genetic scenario, we sought to create two HHT inducible <u>e</u>ndothelial <u>c</u>ell-specific loss <u>o</u>f <u>h</u>eterozygosity (*HHT-*iEC-LOH) models utilizing previously developed mouse models (details described in Methods). Contrary to the well-studied *HHT-iECKO* mice, *HHT-*iEC-LOH mice are born heterozygous (like patients), and deletion of the second gene copy is induced postnatally to mimic the somatic second hit mutation of the remaining wildtype allele (Fig 1a).

**Fig. 1.**
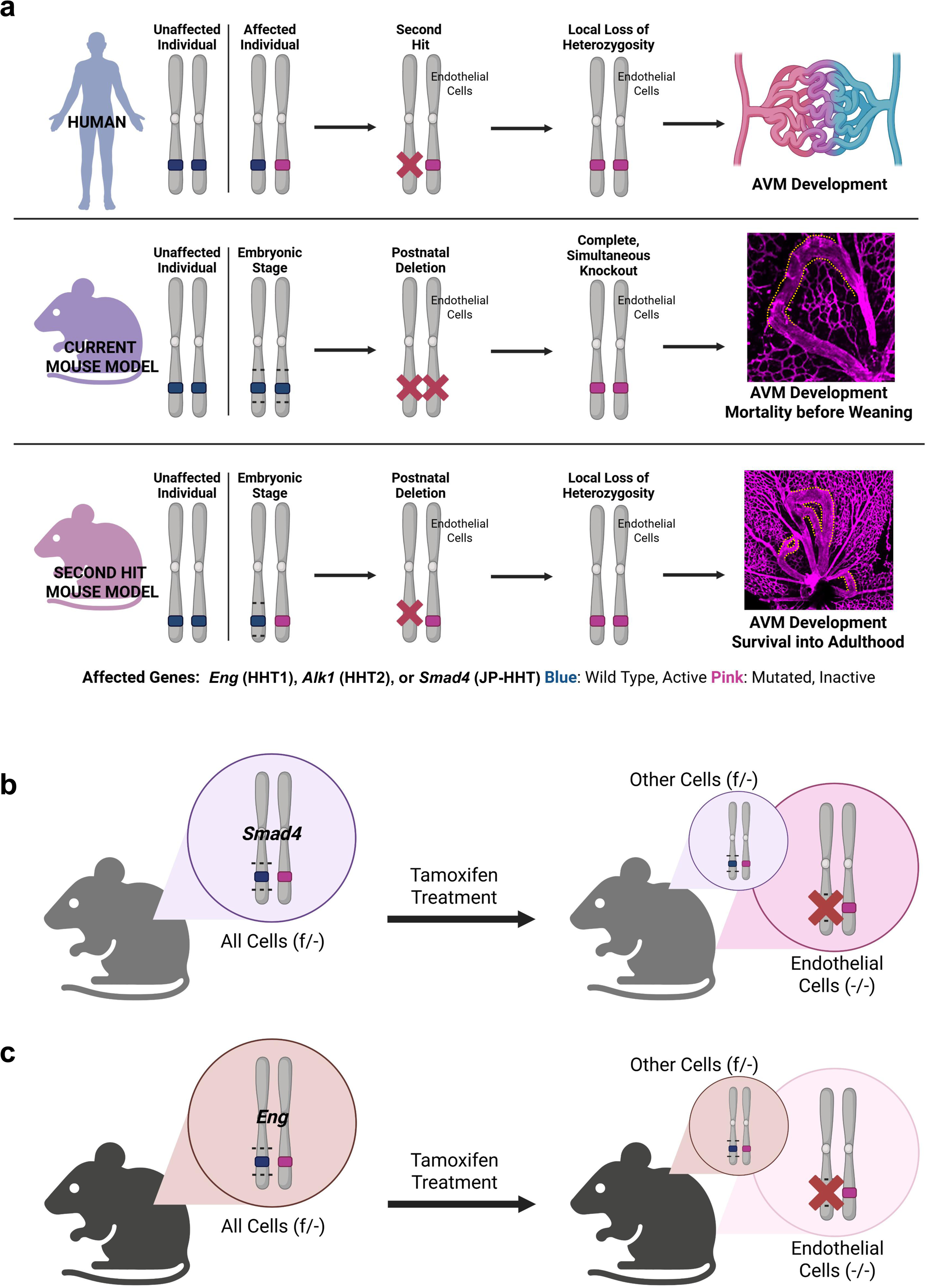
Genetic Rationale and Experimental Framework for Modeling Mosaic Somatic Mutations in HHT. **a.** Comparative graphic noting the differences between HHT patients and the current *HHT-iECKO* models, while highlighting the similarities of the *HHT-*iEC-LOH models to patients’ conditions. People affected with HHT inherit monoallelic inactivating mutations, which makes them heterozygous for expression of SMAD4 or ENG variants. Second hits, such as somatic mutations, can result in localized loss of function (LOH) and contribute toward malformation development. Current HHT inducible EC-specific knock out (*HHT*-iECKO) models do not replicate this genetic progression: mice are born with intact gene expression, and homozygous deletion is induced postnatally. With early gene deletion using standard strategies, mice have been shown to die within the first week or so of life. In contrast, the *HHT*-iEC-LOH mice are born heterozygous (like patients), and deletion of the second gene copy is induced postnatally to create the second somatic mutations. **b.** Graphic representation of the *Smad4-*iEC-LOH model. *Smad4-*iEC-LOH cells start off as heterozygous for active *Smad4*. With tamoxifen treatment, the second copy of *Smad4* is inactivated in some ECs creating localized LOH. *Smad4* is expressed ubiquitously, so heterozygous loss may impact multiple cell types. **c.** Graphic representation of the *Eng-*iEC-LOH model. *Eng-*iEC-LOH cells start off as heterozygous for active *Eng*. With tamoxifen treatment, the second copy of *Eng* is inactivated in some ECs creating localized LOH. *Eng* is also expressed in immune cells, so heterozygous loss may impact these cell types.

To model the genetic mosaicism and distinct pathogenic mechanisms of HHT, we generated and characterized two iEC-LOH models: *Smad4* and *Eng*. *Smad4* was generated and characterized first, with the assumption that heterozygous *Smad4* activity in non-ECs may impact the phenotype, given its ubiquitous expression[55] (Fig 1b). Subsequently, the *Eng model* was generated and characterized, representing a more prominent variant of HHT. *Eng* is also actively expressed in non-ECs, such as immune cells[56], suggesting its heterozygous expression could similarly influence the overall phenotype (Fig 1c). The heterozygous background of both models recapitulates patient genotype, while the low-level, EC-specific induction of LOH in *Smad4* or *Eng* replicates the random somatic mutations found in patient malformations.

In order to establish the iEC-LOH models, *Smad4* and *Eng* homozygous conditional flanking loxP (*HHT*^f/f^) mice were first crossed with epiblast-specific *Sox2*-Cre mice to generate germline, heterozygous knockout mice (*HHT*^f/wt^;*Sox2-Cre* or *HHT^-^*^/wt^;*Sox2-Cre)* (Sup Fig 1a,b). These mice were then bred to corresponding conditional floxed mice (*HHT*^f/f^) to produce *HHT*^f/-^ mice lacking *Sox2-Cre* (Sup Fig 1b), thereby establishing the heterozygous loss of function genetic background observed in HHT patients. Next, *HHT*^f/-^ mice were crossed with *HHT^f/f^;Cdh5-Cre^ERT2^* mice, which contain the tamoxifen inducible <u>EC</u>-specific *Cre* driver Cdh5 (Sup Fig 1c,d). This particular mating strategy results in the following genotypes, which will be compared throughout the studies: *HHT*^f/-^, *HHT*^f/f^, *HHT^f/f^;Cdh5-Cre^ERT2^* (referrred to as *HHT-iECKO*) and *HHT^f/-^;Cdh5-Cre^ERT2^* (referred to as *HHT*-iEC-LOH). The established *HHT-iECKO* model features wildtype non-ECs and inducible biallelic loss in ECs. In contrast, the novel *HHT*-iEC-LOH mice are comprised of all heterozygous cells (the *HHT^f/-^* background) and biallelic loss in ECs upon tamoxifen induction. The remaining control groups, *HHT^f/-^* (heterozygous loss in all cells only) and *HHT^f/f^*(true wild-type), will be used for comparison.

### Partial biallelic loss is sufficient to cause AVM formation in a heterozygous background

Multiple sequencing studies of HHT patient telangiectasia and AVM tissue have revealed the mosaic nature of lesion development, driven by low frequency somatic mutations. These inactivating mutations in the corresponding wildtype allele led to biallelic loss of function in a subset of ECs[22–26]. To determine if our *HHT*-iEC-LOH models could recapitulate this mosaicism, we incorporated the *ROSA mTmG* Cre-reporter line into the LOH models (*HHT^f/-^;mTmG^+/wt^;Cdh5-Cre^ERT2^;* referred to as *HHT*-iEC-LOH;*mTmG*) (Sup Fig 1e). The *mTmG* reporter system allows expression of *tdTomato* in all cells. However, in the presence of Cre-recombinase, *tdTomato* expression is turned off and *GFP* is subsequently turned on, allowing heterozygous loss of function ECs (*tdTomato* positive, non-LOH ECs) to be distinguished from those harboring the induced gene deletion, thereby mimicking the somatic mutation (GFP positive, LOH ECs with biallelic loss of function)[50]. To induce mosaic recombination, low doses of tamoxifen were administered at P1 to establish the minimal dose required to induce retinal AVM formation (Fig 2a,b and data not shown). The retinal vasculature was examined because it is a prominent and established model system for HHT-related AVM studies, particularly those utilizing iECKO models. Based on established *Smad4*- and *Eng-iECKO* studies, we assessed retinas at P7, finding that minimal doses of 5 μg and 2.5 μg of tamoxifen were required to promote consistent AVM formation in the *Smad4*- and *Eng*-iEC-LOH;*mTmG* models, respectively (Fig 2a,b).

**Fig. 2.**
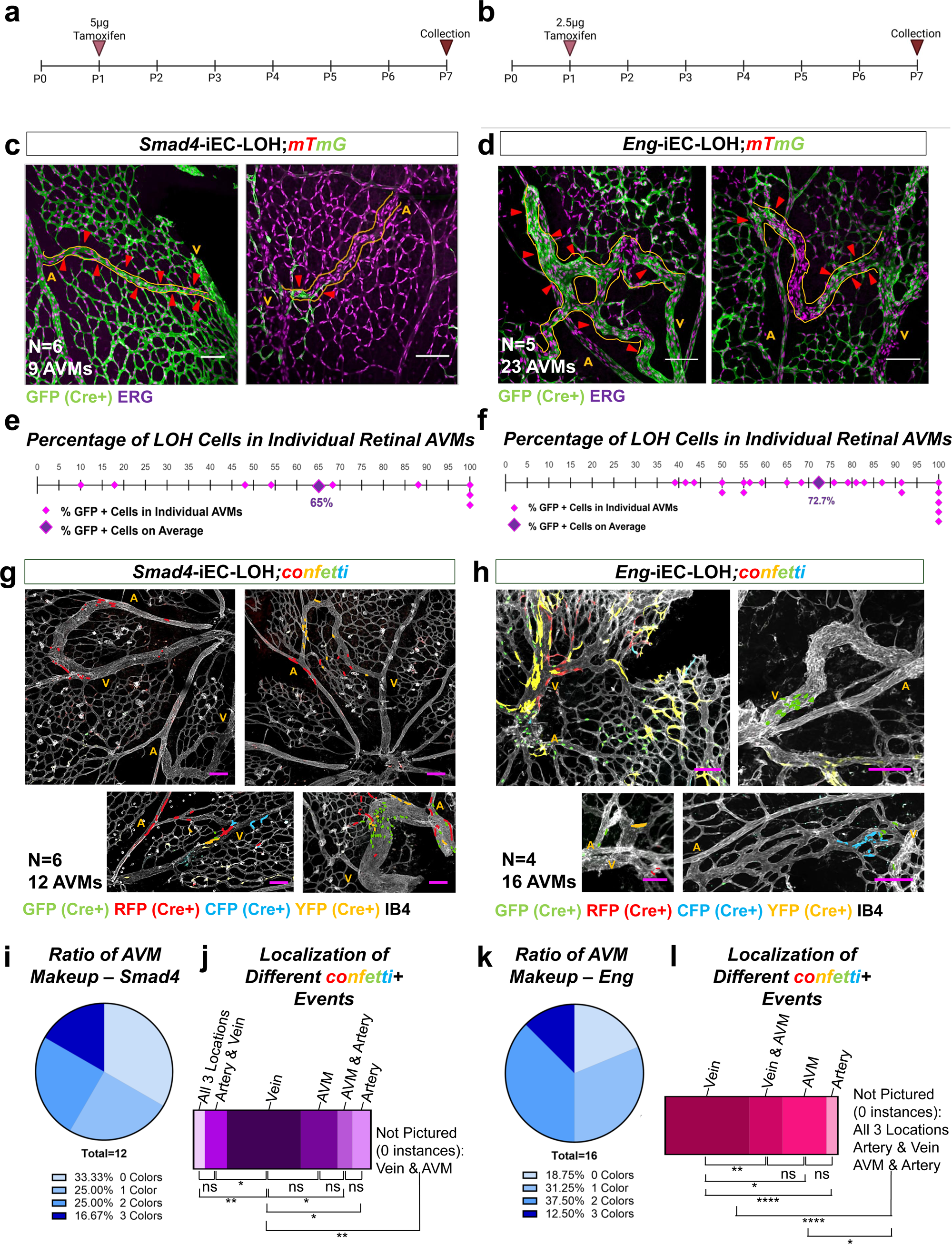
*HHT*-iEC-LOH mice mimic patient mosaicism in the retina and showcase multiple, distinct LOH events contribute to the development of individual AVMs. **a,b.** Tamoxifen treatment strategy for *Smad4 and Eng* litters. **c,d.** Immunofluorescent images of *Smad4- and Eng-*iEC-LOH;*mTmG* retinas at P7 with nuclear ERG staining and endogenous eGFP, which marks the LOH recombinant ECs. Red arrowheads denote areas of GFP positive ECs within the AVM. A, artery; V, vein. Scale bars = 100µm. **e,f.** Quantification of the eGFP composition within AVMs of *Smad4- and Eng-*iEC-LOH;*mTmG* retinas. Pink diamonds represent individual AVMs, while the purple diamond represents the average. **g,h.** Immunofluorescent images of *Smad4- and Eng-*iEC-LOH;*confetti* retinas at P7 with Isolectin B4 (IB4) staining and the endogenous reporters (GFP, RFP, CFP and YFP). Reporter colors highlighted for ease of view. Scale bars = 100µm. **i,k.** Pie graphs depicting the ratio of the AVM makeup in Smad4- (i) and Eng-iEC-LOH (k) retinas based on the number of colors within the AVM. **j,l.** Localization of reporter positive ECs in Smad4- (j) and Eng-iEC-LOH (l) AVMs. For j and l, levels were compared using an Ordinary One-Way ANOVA with Fisher’s LSD. ns = not significant; * = P-value<0.05; ** = P-value<0.01; **** = P-value<0.0001.

To assess AVM mosaicism, *Smad4*- and *Eng*-iEC-LOH;*mTmG* retinas were immunofluorescently labeled for the EC nuclear marker ERG. Using this strategy, all ECs were identified by ERG expression, allowing us to distinguish LOH (GFP+, tdTomato-) from non-LOH (GFP-, tdTomato+) ECs within AVMs. For this study, retinal AVMs were identified as direct shunts between arteries and veins, with receding capillary beds, and with larger diameters than the surrounding capillary bed. Analyzing 9 total AVMs across 6 *Smad4-*iEC-LOH;*mTmG* retinas at P7, the AVMs displayed a wide-ranging composition of LOH versus non-LOH ECs (Fig 2c,e). While some AVMs were composed of as little as 10% of LOH ECs, multiple appeared to be completely GFP, with the average LOH composition being 65% (Fig 2e). In *Eng-*iEC-LOH;*mTmG* mice, 25 total AVMs from 5 different retinas were analyzed, showing a similar broad range from approximately 40% LOH ECs up to 100% (Fig 2d,f). On average, *Eng*-iEC-LOH AVMs were composed of 73% LOH ECs. In both models, most AVMs were a mixture of LOH and non-LOH ECs (6/9 in *Smad4*-iEC-LOH; 18/23 in *Eng*-iEC-LOH). These results closely resemble the mosaic human patient data, confirming the utility of these models to recapitulate and study the focal biallelic loss of function associated with lesion development. Additionally, the efficiency of GFP as a reporter for genetic loss was confirmed by immunostaining *Eng-*iEC-LOH;*mTmG* retinas for ENG. All samples showed less than 3% colocalization of ENG staining with GFP expression, indicating a strong correlation between genetic loss of *Eng* (lack of expression) and GFP expression marking LOH ECs (Sup Fig 3). Performing the same experiment for Smad4 is difficult given its ubiquitous expression; however, we note that the loxP sites flanking exon 8 of *Smad4* (Sup Fig 2a) and the *tdTomato* gene in *mTmG* are nearly equidistant (approximately 1kb), which may be indicative of effectively marking the Smad4-(GFP+) LOH ECs.

### Multiple, distinct second hits contribute towards the development of individual AVMs

Although the mTmG reporter enabled us to assess the level of mosaicism in each lesion, it did not provide a means to determine whether the LOH ECs within the heterogeneous lesion derived from single or multiple LOH events. To address this distinction, we crossed *HHT*-iEC-LOH mice with the ROSA confetti reporter line (*HHT^f/-^;confetti^+/wt^;Cdh5-Cre^ERT2^;* referred to as *HHT*-iEC-LOH;confetti) (Sup Fig 1f). The confetti reporter system initially exhibits no reporter expression. However, upon tamoxifen induction, *Cre*-recombinase stochastically activates one of four distinct fluorescent reporters (GFP, RFP, CFP and YFP), and that specific reporter expression is subsequently maintained in all daughter cells of the recombined lineage. Retinas from *HHT*-iEC-LOH;confetti mice were immunolabeled with Isolectin B4 (IB4) in far red to mark all ECs, and distinct local expansions within each AVM were characterized based on the different fluorescent markers. It should be noted that due to chance, multiple clonal expansion events can occur with the same random reporter color[51]. Because these instances cannot be definitively distinguished, we conservatively identified single mutation events as a single color. Therefore, even if the same color appeared in multiple distinct locales within an AVM, it was recorded as a single event, though these may theoretically represent independent mutations. In addition, because the confetti reporter system is equally likely to result in no reporter color presence upon *Cre* induction, it is not possible to capture the full composition of LOH ECs within an AVM, unlike the *mTmG* reporter.

To analyze the confetti events, we investigated the distribution of fluorescent ECs from 12 *Smad4*-iEC-LOH;*confetti* AVMs (6 retinas) and 16 *Eng*-iEC-LOH;*confetti* AVMs (4 retinas) (Fig 2g,h). We first categorized the number of distinct EC color populations (lineages) within each lesion (Fig 2i,k). *Smad4*-iEC-LOH;confetti AVMs comprised lesions ranging from no recombination events (no color; 33.33%) up to 3 distinct lineages: 1 color (25%), 2 colors (25%) and 3 colors (16.67%) (Fig 2I). *Eng*-iEC-LOH;*confetti* AVMs exhibited a similar distribution of lineages but at different rates: no color (18.75%), 1 color (31.25%), 2 colors (37.5%), and 3 colors (12.50%) (Fig 2k). Comparatively, Smad4 AVMs were most frequently observed with no reporter signal (33.33%), while Eng AVMs were most frequently observed with 2 distinct lineages (37.5%). Moreover, 50% of the Smad4 AVMs consisted of 1 or 2 colors/lineages, compared to 68.75% of Eng AVMs. In both models, 3 colors were the least observed number of lineages. These results suggest that more LOH recombination events are required to initiate AVM development in the *Eng-*iEC-LOH model compared to the *Smad4-*iEC-LOH model, a finding that correlates with the higher average LOH EC composition observed in *Eng-*iEC-LOH;*mTmG* AVMs (Fig 2c-f). Furthermore, given that the *mTmG* lineage analysis indicated that at least one recombination event must be present in an AVM, we infer that those AVMs with no confetti signal must harbor at least one LOH recombination event that was not captured by the stochastic confetti system. Taking this into consideration, the data strongly suggests that AVMs from either the Smad4 or Eng LOH models arise from at least one clonal expansion but are typically driven by the contribution of several independent LOH recombination events.

Furthermore, the spatial localization of different confetti events was calculated for individual AVMs based on whether a single color/lineage was found only on or near the vein side, on or near the artery side, distributed within the AVM body, or as a combination of these regions. A normalized localization ratio was then assigned to each separate AVM, ensuring that lesions with more distinct confetti events were not weighted more heavily than those with fewer. For example, if only one lineage was observed within an AVM and found exclusively on the vein side, it was assigned a 100 ratio for the “Vein” localization, as 100% of the observed lineages in that AVM were found there. Similarly, if three colors were observed within the same AVM and were all observed on the vein side, a 100 ratio would be assigned to “Vein”. However, if two distinct lineages were observed with different localization patterns—such as one only on the vein side and one only on the artery side—a 50 ratio was assigned to the “Vein” and “Artery” categories separately, reflecting that 50% of the AVM’s lineages were found at each location. For every AVM, the resulting localization ratios across all categories should sum to 100. These normalized datasets were then used to compare the significance of localization ratios, with the bar graphs representing the average ratio of each localization pattern generated by averaging the individual AVM ratios (i.e., dividing the sum of ratios for each localization pattern by the total number of AVMs analyzed).

Based on the spatial analysis of confetti events, most recombined lineages observed in the Smad4-based AVMs are localized to the venous side of the AVM when compared to other regional patterns, except when compared to the AVM body (Fig 2j). Similarly, the majority of recombined lineages in Eng-LOH AVMs were localized only to the venous side compared to any other regional pattern (Fig 2l). Interestingly, compared to S*mad4*-iEC-LOH;*confetti* AVMs, which showed distinct lineages spanning multiple different regions (i.e., artery and vein; AVM body and vein; all three regions), confetti events were largely confined to only one region of the vasculature in Eng deficient AVMs, with the exception of the AVM body-vein combination. Nonetheless, the overall trend of lineage localization to the venous side of the AVMs highlights the possible importance of vein dysregulation in lesion pathogenesis.

### *HHT-*iEC-LOH mice develop retinal vasculature defects with more consistency and severity than the *HHT-iECKO* mice

After analyzing the LOH/non-LOH cellular makeup of the *HHT-*iEC-LOH AVMs, we performed a detailed characterization of the retinal vascular defects. The postnatal mouse retina is a robust model to observe vascular defects due to its conserved development and consistent alternating patterning of arteries and veins connected by capillary beds[54]. Further, this is the most common tissue analyzed with *HHT*-*iECKO* mice. Based on our mating strategy, the following genotypes would be equally present in the litters allowing us to compare the LOH and iECKO models in addition to their control counterparts (*HHT*^f/-^ and *HHT*^f/f^). Using the same induction regimen with low doses of tamoxifen (Fig 2a,b), we assessed AVM development and vessel calibers (vessel enlargement in *HHT*-*iECKO* mice is a well-defined characteristic of these models) in P7 retinas. Analysis of the Smad4 models treated with 5µg of tamoxifen and stained for IB4 and ERG showed clear distinctions. Retinas of *Smad4-*iEC-LOH mice displayed consistent, robust AVM formation (Fig 3a; Sup Fig 4). Approximately 60% of *Smad4*-iEC-LOH retinas developed at least one AVM, which was significantly higher than the prevalence of *Smad4-iECKO* retinal AVMs (about 15%) (Fig 3c). Moreover, *Smad4*-iEC-LOH mice displayed significantly more AVMs per retina on average than *Smad4-iECKO* mice and controls (Fig 3d). It was also noted that LOH AVMs were more robust and easier to identify than the iECKO AVMs. Vessel diameter was also measured to explore other vascular defects associated with HHT. Artery and vein calibers of *Smad4*-iEC-LOH mice were significantly larger than those of *Smad4-iECKO* mice and respective controls (Fig 3e,f). Interestingly, *Smad4-iECKO* artery and veins sizes were comparable to those in the *Smad4^f/-^* mice with both being significantly bigger than *Smad4^f/f^* retinas (Fig 3 e,f). This result suggests that in terms of vessel enlargement, a heterozygous background, even in the absence of LOH induction (i.e., somatic mutation), exhibits a significant vascular defect as well. Therefore, the *Smad4*-iEC-LOH model may provide a significantly more penetrant and robust platform for studying HHT pathogenesis compared to the conventional iECKO model.

**Fig. 3.**
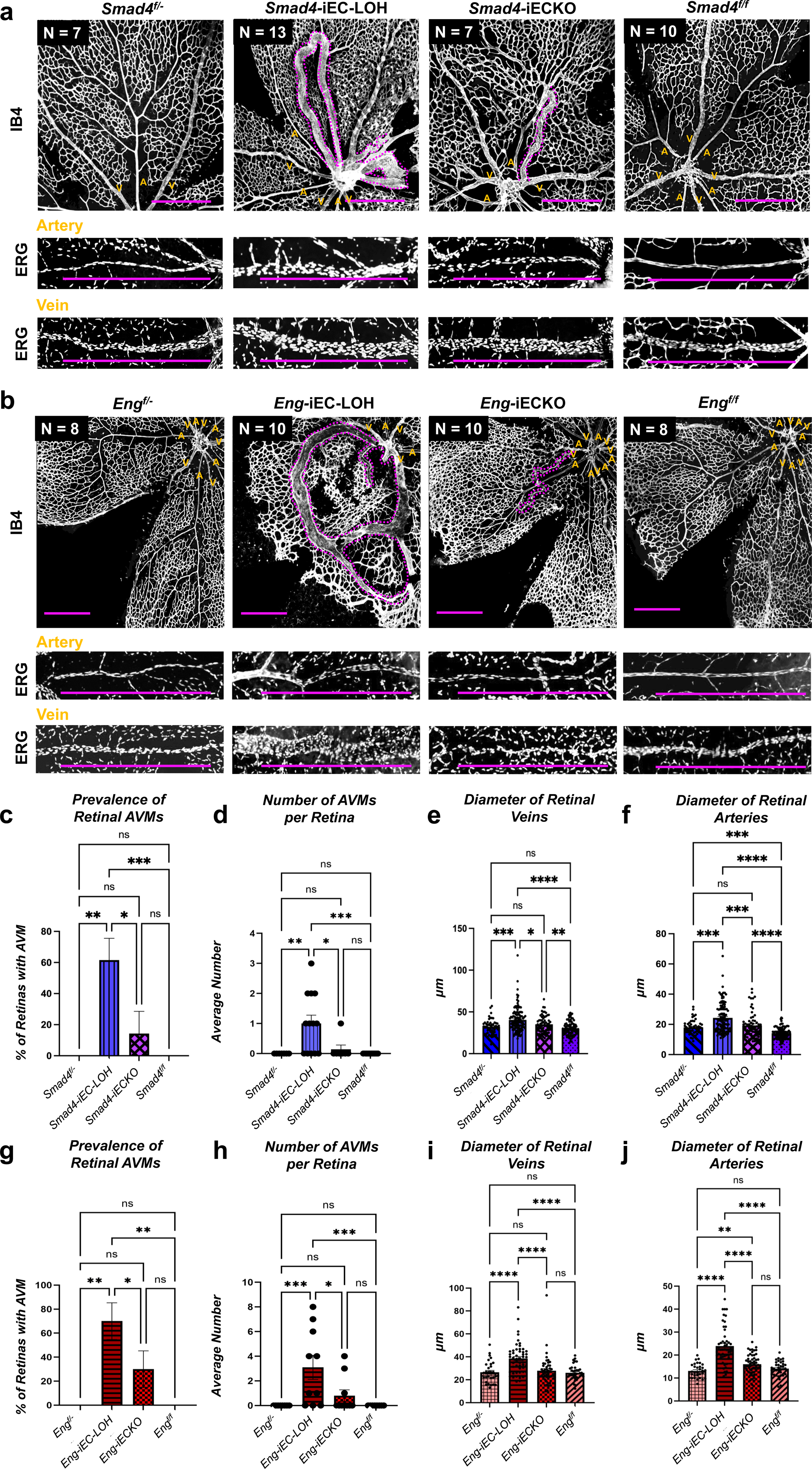
Enhanced AVM development and increased vessel diameters in *HHT-*iEC-LOH retinas compared to *HHT*-iECKO mice. **a,b.** Retinal images of all *Smad4 and Eng* models (LOH, iECKO, f/- and f/f) immunostained with Isolectin B4 (IB4) or ERG at P7. AVMs are outlined in purple dotted lines. Closeup images of arteries and veins are also presented. All scale bars = 500µm. A, artery; V, vein. **c.** Prevalence of AVMs in the *Smad4* models showed an increase of retinas with AVMs in the *Smad4-*iEC-LOH mice when compared to all other genotypes. **d.** The average number of AVMs per retina is higher in *Smad4-*iEC-LOH mice than all other genotypes. **e.** Retinal vein diameters in *Smad4-*iEC-LOH mice are higher than all other genotypes. *Smad4-iECKO* mice have more dilated veins than *Smad4^f/f^* controls but are comparable to *Smad4^f/-^* heterozygous mice. **f.** Retinal artery diameters in *Smad4-*iEC-LOH mice are higher than all other genotypes. *Smad4-iECKO* mice have more dilated arteries than *Smad4^f/f^*controls but are comparable to *Smad4^f/-^* mice. *Smad4^f/-^*retinas also exhibit increased arterial diameters when compared to *Smad4^f/f^*controls. **g.** Prevalence of retinal AVMs in the *Eng* models showed an increase of retinas with AVMs in the *Eng-*iEC-LOH mice when compared to all other genotypes. **h.** The average number of AVMs per retina is higher in *Eng-*iEC-LOH mice than all other genotypes. **i.** Retinal vein diameters in *Eng-*iEC-LOH mice are higher than all other genotypes. **j.** Retinal artery diameters in *Eng-*iEC-LOH mice are higher than all other genotypes. *Eng-iECKO* mice have more dilated arteries than *Eng^f/-^*retinas. Vessel diameters were measured using ImageJ. For c–j, comparisons across groups were performed using a Kruskal-Wallis test with Dunn’s post-hoc. ns = not significant; * = P-value<0.05; ** = P-value<0.01; *** = P-value<0.001; **** = P-value<0.0001. Error bars represent□±□SEM.

We next utilized the same approach to investigate the *Eng*-iEC-LOH model. *Eng-*iEC-LOH pups given 2.5µg of tamoxifen at P1 and collected at P7 displayed comparable outcomes. Retinas of *Eng-*iEC-LOH mice displayed consistent, robust AVM formation and severe vascular defects (Fig 3b; Sup Fig 4). Approximately 70% of *Eng-*iEC-LOH retinas developed at least one AVM, which was significantly higher than the prevalence of *Eng iECKO* retinal AVMs (Fig 3g). Furthermore, *Eng-*iEC-LOH mice displayed significantly more AVMs per retina on average than *Eng-iECKO* mice and controls (Fig 3h). Similar to the Smad4 studies, AVMs in *Eng-*iEC-LOH were more substantial and much easier to identify than those in *Eng-iECKO* mice. However, in comparison to Smad4-LOH AVMs, AVMs in *Eng-*iEC-LOH retinas formed at a much greater rate (Fig 3d versus h). Assessment of arteries and veins showed that these vessels were significantly more dilated in *Eng-*iEC-LOH mice than those of *Eng-iECKO* mice and respective controls (Fig 3i,j). Interestingly, *Eng-iECKO* artery and veins sizes were comparable to those in the *Eng^f/f^* mice (Fig 3i,j), while vein diameters were also comparable to *Eng^f/-^* mice (Fig 3i). In the context of both LOH models, *Eng-*iEC-LOH mice displayed worse retinal vasculature phenotypes than *Smad4-*iEC-LOH mice, with higher retinal AVM prevalence, more AVMs per retina, and more exaggerated differences in vessel diameter when compared to controls. All together, these observations demonstrated the significant pathological severity driven by the LOH models, highlighting that even subsets of LOH-induced ECs are sufficient to drive substantial vascular morphological changes.

### Cerebrovascular abnormalities are prevalent in the *HHT*-iEC-LOH mice

While the retina is a highly beneficial model system, the brain has more physiological significance since the occurrence of brain AVMs in HHT patients is more prevalent than retinal AVMs, which are rare[57–59]. To identify cerebral vascular defects, we used an established method of perfusing the vasculature with a blue latex dye[43]. In a typical vascular network, the latex is unable to flow into capillary beds and is therefore absent in the veins. In a dilated capillary bed (i.e., AVM), latex flows from the arteries into the venous system and is therefore used as a proxy to identify AVMs[39, 40, 43, 60–62] (Fig 4a). For these experiments, four major veins were analyzed for blue latex presence: rostral rhinal veins, anterior cerebral veins, basal veins, and inferior cerebral veins (Fig 4b, Sup Fig 5a); and four major artery diameters were measured: anterior communication artery, middle communicating artery, posterior communicating artery, and basilar artery (Sup Fig 5a). Veins were counted as positive for blue latex presence if at least one of the two corresponding major veins had latex, and arteries located on the right side of the brain were measured to maintain consistency.

**Fig. 4.**
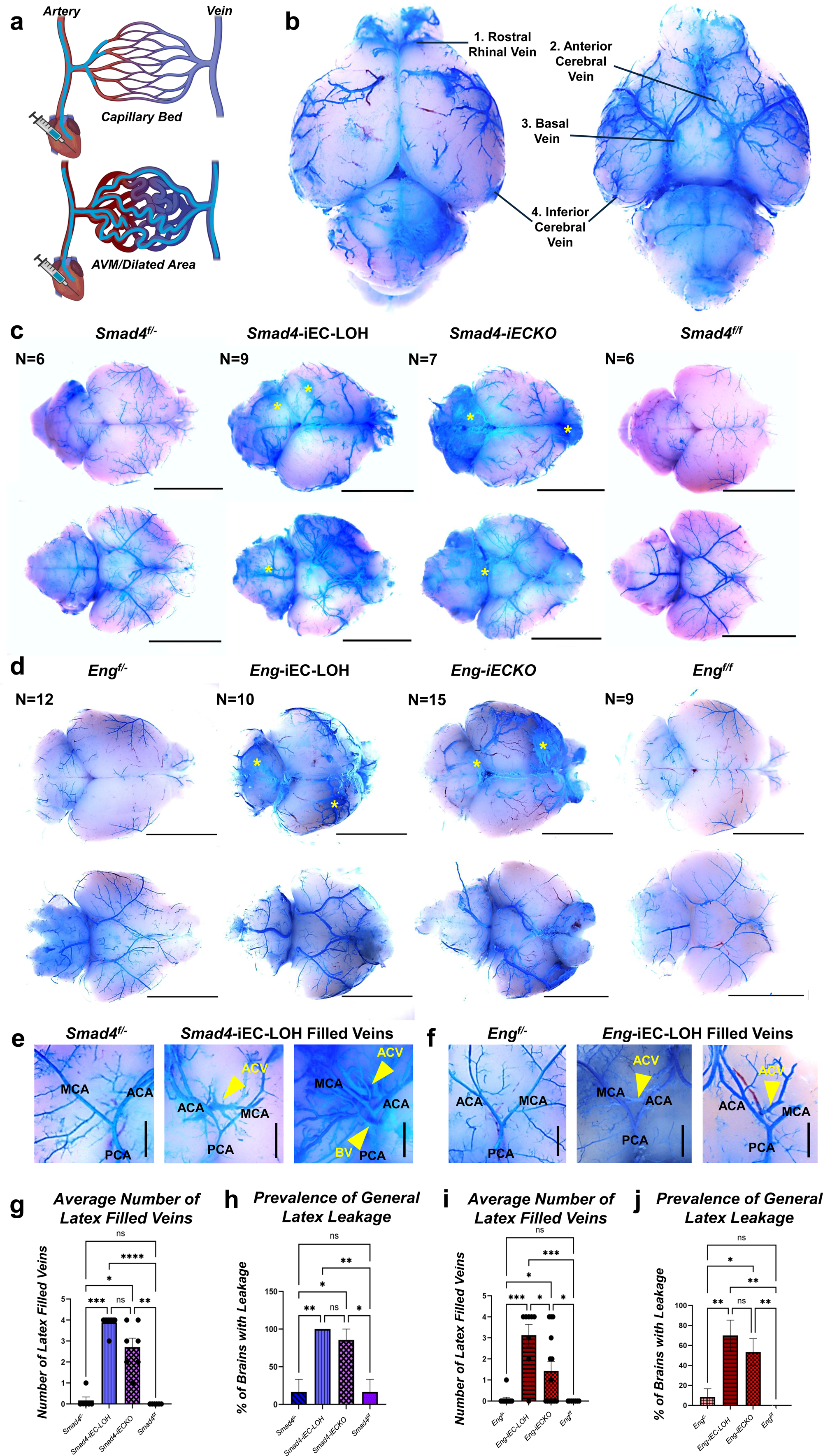
Blue latex perfusion reveals enhanced cerebrovascular AVM penetrance and arterial dilation in *HHT-*iEC-LOH models. **a.** Schematic of blue latex vasculature perfusions. In a normal capillary bed, blue latex is impeded and remains in the arterial system. In dilated capillaries, blue latex is able to flow into the venous system, indicative of the presence of AVMs. **b.** Diagram of the four major veins characterized for blue latex presence (*Smad4-*iEC-LOH brain is shown): rostral rhinal vein, anterior cerebral vein (ACV), basal vein (BV), and inferior cerebral vein. **c, d.** Bright field imaging of *Smad4* and *Eng* brains perfused with blue latex. Dorsal (top) and ventral (bottom) views are shown. Yellow asterisk = latex leakage. Scale bars = 5000 µm. **e,f.** Representative close-up images of latex-filled veins in *Smad4-* and *Eng-*iEC-LOH and corresponding control f/-brains. Scale bars = 1000 µm. Yellow arrowheads point at veins filled with latex. ACA, anterior communicating artery; MCA, middle communicating artery; PCA, posterior communicating artery. **g.** The average number of latex-filled veins is comparable in *Smad4-*iEC-LOH and *Smad4-iECKO* brains, which are both higher than controls. **h.** Prevalence of general latex leakage is comparable in *Smad4-*iEC-LOH and *Smad4-iECKO* mice, which are both higher than controls. **i.** The average number of latex -veins is higher in *Eng-*iEC-LOH compared to all other genotypes. **j.** The prevalence of general latex leakage is comparable in *Eng-*iEC-LOH and Eng*-iECKO* mice, which are both higher than controls. For g-j, comparisons were performed using a Kruskal-Wallis test with Dunn’s post-hoc. ns = not significant; * = P-value<0.05; ** = P-value<0.01; *** = P-value<0.001; **** = P-value<0.0001. Error bars represent□±□SEM.

Grossly, brains of the low dose, tamoxifen treated *Smad4-*iEC-LOH mice exhibited comparable, and sometimes worse, vasculature defects when compared to *Smad4-iECKO* mice, though the defects in both mutant Smad4 genotypes were distinct compared to the control brains. (Fig 4c). On average, both Smad4 mutant genotypes had comparable numbers of veins filled with latex (Fig 4c,e,g), which is indicative of localized dilated capillary beds (AVMs). Presence of latex filled veins was prominently observed in the anterior cerebral vein and basal vein of the mutant brains (Fig 4e). Specifically, however, *Smad4-*iEC-LOH brains had more anterior and inferior cerebral veins filled with latex than *Smad4-iECKO* mice (Sup Fig 5c,e). In addition, the basilar artery and posterior communicating artery were significantly more dilated in *Smad4-*iEC-LOH mice than controls (Sup Fig 5h,i), but no significant increases in dilation were observed compared to *Smad4-iECKO* mice (Sup Fig 5h,i). Furthermore, generalized leakage of latex (asterisks) outside of the vasculature was observed in both the Smad4 LOH and iECKO models, indicating severe vessel compromise, while dye was retained in the arteries of both control brains (Fig 4h). Thus, while AVM prevalence was similar between the two Smad4 mutant models, the *Smad4-*iEC-LOH mice exhibited a more severe vascular compromise, characterized by greater arterial dilation and dye leakage.

Analysis of the Eng models given 2.5µg of tamoxifen at P1 revealed that P7 *Eng-*iEC-LOH brains displayed similar overall vascular defects when compared grossly to *Eng-iECKO* mice (Fig 4d). However, upon quantification, *Eng-*iEC-LOH mice had more veins filled with latex, while neither control genotypes exhibited a significant presence of latex-filled vessels (Fig 4f,i). *Eng-*iEC-LOH brains had more rostral rhinal, basal, and inferior cerebral veins filled with latex than *Eng-iECKO* mice (Sup Fig 5j,l,m). Further, the basilar artery was significantly more dilated in *Eng-*iEC-LOH brains than all other groups (Sup Fig 5p). The middle communicating, anterior communicating, and posterior communicating arteries were all more dilated in *Eng-*iEC-LOH mice when compared to both control groups (Sup Fig 5n,o,q), whereas the middle communicating and basal arteries of the *Eng-iECKO* brains were only significantly increased when compared to the *Smad4*^f/-^ controls alone (Sup Fig 5n,p). In terms of latex leakage, Eng LOH and *Eng-iECKO* mice showed comparable leakage, but significantly higher when compared to control mice (Fig 4d,j). Together, these data demonstrate that the *Eng-*iEC-LOH model generates a cerebral AVM phenotype with greater penetrance and vascular compromise compared to the *Eng-iECKO* model, mirroring the enhanced pathology observed in the retina.

Comparatively, the *Smad4-*iEC-LOH mice had a higher average number of veins filled with latex and higher prevalence of generalized leakage than the *Eng-*iEC-LOH mice. Diameter changes in the *Smad4-*iEC-LOH brains were localized to the vessels near the posterior aspect of the Circle of Willis, while the *Eng-*iEC-LOH mice had increased artery diameters around the entire Circle of Willis. In the brain, *Smad4-iECKO* mice also showed a more comparable phenotype to the *Smad4-*iEC-LOH mice, while *Eng-*iEC-LOH mice maintained a generally worse phenotype than *Eng-iECKO* mice.

### Red blood cell leakage in *HHT-*iEC-LOH retinas

Bleeding in HHT can arise due to ruptured vessels or disruptions in cell connections, leading to reduced vessel integrity and increased permeability[63, 64]. While it is well-established that those afflicted with HHT experience bleeding in the nose, gastrointestinal tract, and from telangiectasias[65–67], there is limited characterization of bleeding and vessel leakiness in mouse models of HHT. To investigate potential blood retinal barrier defects at P7, TER119 immunostaining was performed following clearance of the vessels with PBS to identify red blood cell (RBC) presence in the perivascular space[68]. Using IB4 to identify the blood vessels and TER119 to mark RBCs, we identified a significant presence of RBCs, as measured by coverage area, in *Smad4-*iEC-LOH retinas compared to all other groups (Fig 5a,c). RBC leakage in the *Smad4*-iEC-LOH models was significantly higher than the *Smad4-iECKO* retinas, with the latter showing a level of RBC leakiness above the control groups, which was minimal. The RBC leakage was observed around areas of hemorrhaging, as well as around relatively “normal” capillary beds (Fig 5b). In a similar fashion, staining for RBCs also revealed a significant increase of RBC coverage area in *Eng-*iEC-LOH retinas compared to controls, but not when compared to *Eng-iECKO* retinas (Fig 5d,f). Like the Smad-LOH retinas, RBC leakage was observed around areas of AVMs, as well as around relatively “normal” capillary beds (Fig 5e). Overall, these experiments demonstrate a higher level of compromised vessel integrity and subsequent bleeding in the LOH model, while also showing that RBC leakage from the blood vessels is not just limited to hemorrhaging, but also from AVMs and even the vascular beds which appear relatively normal morphologically. When comparing both LOH models, the *Smad4-*iEC-LOH mice exhibited much higher RBC coverage outside of the vasculature than *Eng-*iEC-LOH retinas (∼1×10^6^ µm^2^ vs ∼1×10^5^ µm^2^ on average), indicating more problems with retinal permeability deficits in the *Smad4-*iEC-LOH mice. This result was similar to the latex dye perfusion experiments where generalized leaking of the dye was more abundant in the brains of *Smad4-*iEC-LOH mice compared to *Eng-*iEC-LOH mice (Fig 4), further indicating increased vessel integrity and permeability deficits in the Smad4 LOH mutants.

**Fig. 5.**
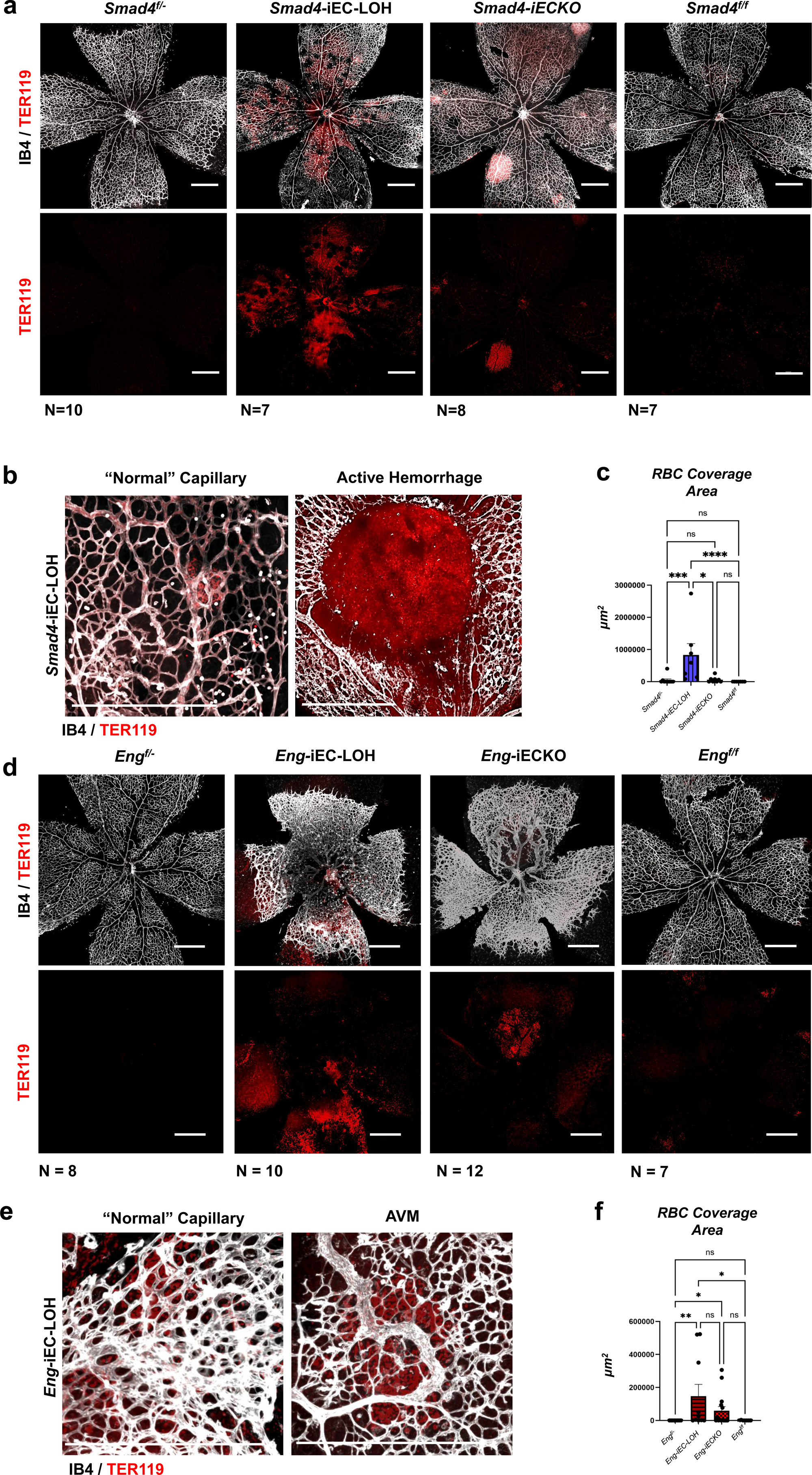
Significant red blood cell leakage is evident in *HHT*-iEC-LOH retinas. **a.** Retinal images of all *Smad4* models immunolabeled with Isolectin B4 (IB4) and TER119 (erythroid lineage / red blood cell (RBC) marker) at P7. Scale bars = 500 µm. Note presence of RBCs outside of the blood vessels, even in areas where AVMs are not present. **b.** Close-up images of *Smad4-*iEC-LOH retinas show RBCs around relatively normal developed capillary networks and at a site of hemorrhage. **c.** RBC coverage outside of the vascular space was measured following clearance of the vessels by PBS perfusion. The coverage area is significantly higher in *Smad4-*iEC-LOH retinas when compared to all other mice. **d.** Retinal images of all *Eng* models immunolabeled with Isolectin B4 (IB4) and TER119. All scale bars = 500 µm. Note presence of RBCs outside of the blood vessels, even in areas where AVMs are not present. **e.** IB4 and TER119 detection in close-up images of *Eng* iEC-LOH retinas reveals the presence of RBCs around AVMs and capillary networks lacking AVMs. **f.** RBC coverage outside of the vascular space was measured following clearance of the vessels by PBS perfusion. The coverage area is significantly higher in *Eng-*iEC-LOH mice when compared to *Eng^f/-^* and control *Eng^f/f^*retinas, while *Eng-iECKO* mice have more RBC coverage than *Eng^f/-^*mice. For c and f, coverage area was measured using ImageJ and compared using a Kruskal-Wallis test with Dunn’s post-hoc. ns = not significant; * = P-value<0.05; ** = P-value<0.01; *** = P-value<0.001; **** = P-value<0.0001. Error bars represent□±□SEM.

### *HHT-*iEC-LOH mice exhibit impaired cerebrovascular integrity

To further interrogate vascular integrity, we explored possible blood brain barrier defects using 10 kDa and 70k Da high-weight fluorescent conjugated tracers. These tracers are representative of larger molecules that would be observed in humans, in contrast to 1 kDa tracers which have been used in previous studies[69] (note: in our experience, high-weight tracers are not able to be visualized in the retina). In a typical setting when performing transcardial perfusions, molecules of this size would not be able to pass through the blood brain barrier. Therefore, following PBS clearance, tracer presence should be absent in the brain unless leakage occurs (Sup Fig. 6).

Upon examination of the Smad4 models, *Smad4*-iEC-LOH mice generally exhibited a higher presence of both 10 kDa and 70k Da tracers when compared to all other genotypes (Fig 6a). Regarding the 10 kDa tracer, significantly more *Smad4-*iEC-LOH brains showed presence of fluorescence when compared to *Smad4-iECKO* and *Smad4^f/-^*mice, and they were approaching significance when compared to *Smad4^f/f^*mice (Fig 6c). However, with the 70 kDa tracer, *Smad4-*iEC-LOH mice showed significantly more prevalence of tracer when compared to all other groups (Fig 6d). Interestingly, presence of both tracers in *Smad4-iECKO* brains was similar to controls (Fig 6c,d) signifying defective vessel integrity was largely relegated to the Smad4 LOH mice.

**Fig. 6.**
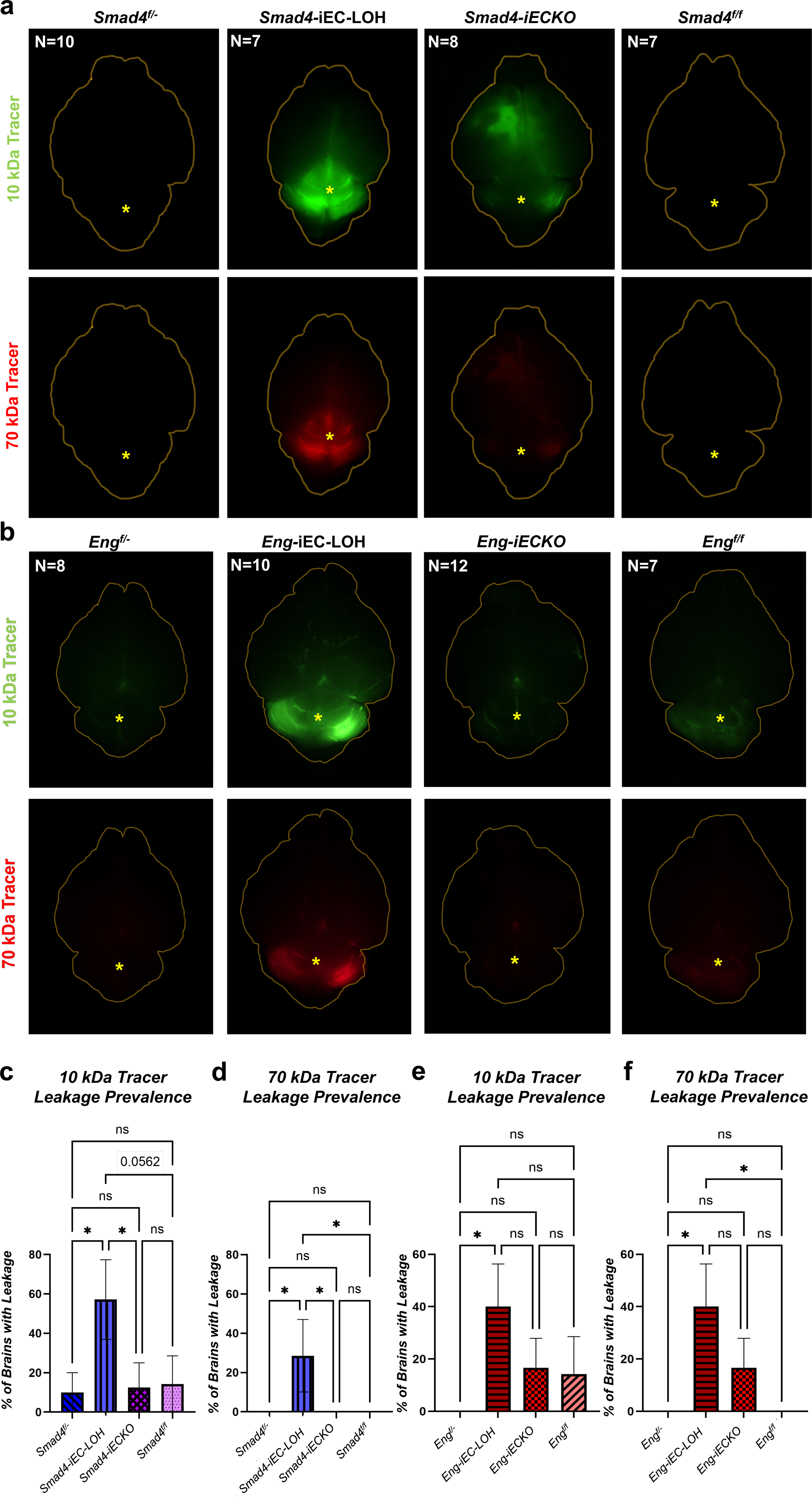
Compromised vascular integrity in *HHT*-iEC-LOH brains visualized by fluorescent dextran extravasation. **a,b.** Fluorescent brain images of *Smad4 and Eng* models perfused with dextran-weighted tracers and cleared with PBS. 10kDA detxran is FITC-conjugated; 70kDa dextran is Texas red-conjugated. Dorsal views are shown and brains are outlined in yellow. Notable leakage of both fluorescent tracers can be seen in the *Smad4- and Eng-*iEC-LOH brains. Tracer leakage was preferentially observed within the cerebellum. **c.** The prevalence of 10kDa tracer leakage is significantly higher in *Smad4-*iEC-LOH than *Smad4-iECKO* and Smad4^f/-^ brains and is trending towards significance when compared to Smad4^f/f^. **d.** The prevalence of 70kDa tracer leakage is significantly higher in *Smad4-*iEC-LOH brains compared to all other genotypes. **e.** The prevalence of 10kDa tracer leakage is significantly higher in *Eng* iEC-LOH than Eng-iECKO, but not more than Engf/- and Engf/f mice. **f.** The prevalence of 70kDa tracer leakage is comparable in *Eng* iEC-LOH and *Eng-iECKO* brains and is significantly higher in *Eng* iEC-LOH than *Eng^f/-^*and *Eng^f/f^*mice. Prevalence was compared using a Kruskal-Wallis test with Dunn’s post-hoc. ns = not significant; * = P-value<0.05. Error bars represent□±□SEM.

Perfused brains of the *Eng-*iEC-LOH mice showed a higher prevalence of both 10 kDa and 70k Da tracers following PBS clearance. However, this difference was only statistically significant for 10kDa when comparing the *Eng-*iEC-LOH mice to *Eng^f/-^* controls (Fig 6e). For 70k Da, *Eng-*iEC-LOH mice had a significantly higher tracer prevalence when compared to both *Eng^f/-^* and *Eng^f/f^* controls (Fig 6f). Thus, unlike the Smad4 models, there was no statistical difference in the amount of tracer detected between the *Eng-*iEC-LOH and *Eng*-iECKO brains.

It is interesting to note that, while 60% of *Smad4-*iEC-LOH brains showed obvious leakage of 10 kDa tracers, only about 30% showed a clear leakage of 70 kDa tracers. In *Eng-*iEC-LOH mice, approximately 40% of brains showed obvious leakage of both 10 kDa and 70 kDa tracers. When comparing the different models, the barrier defects appear more widespread and prevalent in *Smad4-*iEC-LOH mice based on the combined retinal RBC, tracer leakage and latex leakage in the brain. However, the nature of the dysfunction in *Eng-*iEC-LOH mice may represent a more severe or extensive defect, given the larger weight tracers escaped in all instances where tracer leakage was observed. Regarding leakage localization, tracers were frequently detected within the cerebellum in both models (cerebellums marked by asterisks in Fig 6). This suggests that the initial compromise may occur in the vessels supplying the cerebellum.

### Physiological characterization of *Smad4* and *Eng* models

In addition to assessing the vascular defects associated with the different models and corresponding control mice, we also analyzed other characteristics to gain comparative insight into the general health of the mice at P7. Utilizing the same low-dose tamoxifen strategy, we first assessed the sex distribution between the Smad4 and Eng cohorts. Of the seven *Smad4* litters collected, the average litter size was 7.43 pups. The distribution of males and females among these litters was comparable, though trending towards more females (Sup Fig 7a). Of the nine *Endoglin* litters collected, the average litter size was 6.78 pups, and the distribution of males and females among these litters was nearly 50% (Sup Fig 7c).

Next, we measured combined male and female body weights of P7 mice. *Smad4-*iEC-LOH mice had a significantly lower body weight when compared to *Smad4^f/f^* controls (Sup Fig 7b), while the overall average body weight of all genotypes was comparable in the *Eng* litters (Sup Fig 8d). When separating males and females there was no differences in body weight of the different Smad4 male mice (Sup Fig 7e) or between the different Eng groups in both males and females (Sup Fig 7i,j). However, in females, *Smad4^f/-^*pups weighed less than *Smad4^f/f^*, while *Smad4-*iEC-LOH pups weighed significantly less than both the *Smad4^f/f^* and *Smad4-iECKO* mice (Sup Fig 7f). These results suggest that while global growth is largely maintained, the *Smad4*-iEC-LOH genotype specifically impacts neonatal weight gain in a sex-dependent manner.

Additionally, heart weights were measured since they are reported to be larger in *Smad-* and *Eng*-iECKO mice with full gene deletion[44, 70]. With male and female data separated, *Smad4* litters had comparable heart sizes as a ratio of body weight in all genotypes and no differences in body weights within the male and female groupings (Sup Fig 7g,h). Conversely, *Eng*-iEC-LOH males had significantly larger hearts compared to *Eng^f/f^* males (Sup Fig 7k), and *Eng-iECKO* females had significantly larger hearts than *Eng^f/f^* females, though the *Eng-*iEC-LOH trended towards also being bigger (Sup Fig 7l). In summary, while specific genotypes (particularly *Smad4-*iEC-LOH females and *Eng-*iEC-LOH males) exhibit measurable deviations in body and heart weight, these shifts appear to be localized effects rather than an indication of systemic physiological challenges across all models at this developmental stage.

### Low-dose tamoxifen treated HHT-iEC-LOH mice live into adulthood and develop HHT-like pathology

*HHT-iECKO* models with full gene deletion succumb to severe vascular defects upon early Cre induction and fail to live into adulthood, precluding longitudinal studies to monitor AVM development, progression and regression[30, 40, 43]. Since this study utilizes a mosaic recombination strategy, we investigated the ability of these mice to live into adulthood. Of the mice induced at P1 with low doses of tamoxifen, 70% of *Smad4*-iEC-LOH mice and 73% of *Eng-*iEC-LOH mice lived to P70, with minimal or no death being observed in the control groups and even *HHT*-iECKO mice at these specific dosages (Fig 7a,b). We noted that there was a precipitous decline in the health of *HHT*-iEC-LOH after P70 (data not shown). At P70, *HHT-*iEC-LOH mice exhibited noticeable alterations in the cerebral vasculature compared to all other genotypes. Bright-field images of the *HHT*-LOH brains showed signs of enlarged vessels and localized hemorrhages; whereas the blood vasculature of *HHT*-iECKO and control genotypes appeared similarly normal to each other, with only occasional hemorrhages detected in the iECKO mice (Fig 7d,e,h,i).

**Fig. 7.**
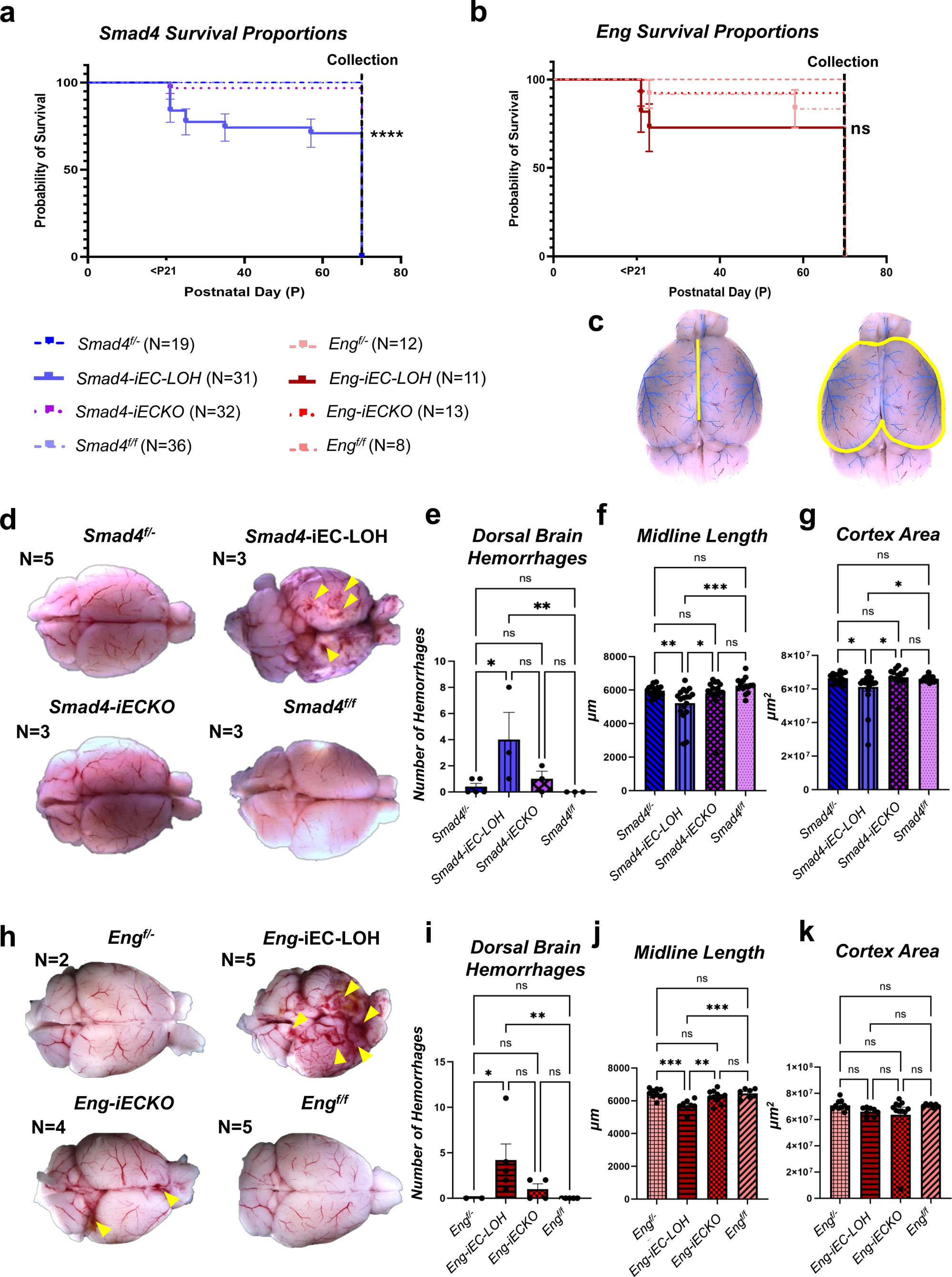
Longitudinal survival and adult brain morphotometry in *HHT*-LOH models. **a**. Survival statistics of *Smad4* models as visualized using a Kaplan – Meir survival curve. One *Smad4-iECKO* mouse died prior to P70, while multiple *Smad4-*iEC-LOH mice died before P70. The P-value for the survival comparison was <0.0001 (error bars represent□±□SEM), showing increased death of the *Smad4-*iEC-LOH mice though survival was near 70% by P70. **b.** Survival statistics of *Eng* models as visualized using a Kaplan – Meir survival curve. Multiple *Eng-iECKO*, *Eng* iEC-LOH, and *Eng^f/-^* mice died before P70. The P-value for the survival comparison was >0.05 (error bars represent□±□SEM), showing similar mortality rates and an approximately 70% survival rate for the Eng LOH mice. **c**. Representative images of midline cortex length (left) and cortical areas (right) in a latex perfused brain. **d**. Bright field images of *Smad4* brains. Yellow arrowheads denote hemorrhages. **e**. More dorsal brain hemorrhages are seen in *Smad* iEC-LOH brains. **f**. The midline length is significantly reduced in *Smad4-*iEC-LOH mice when compared to all other groups. **g**. The cortical area is significantly reduced in *Smad4* iEC-LOH mice when compared to all other groups. **h**. Bright field images of *Eng* brains. Yellow arrowheads highlight hemorrhages. **i**. More dorsal brain hemorrhages are seen in *Eng-*iEC-LOH brains. **j**. The midline length is significantly reduced in *Eng* iEC-LOH mice when compared to all other groups. **k**. The cortical area is unchanged. For **e** and **i**, dorsal brain hemorrhages were compared using a Kruskal-Wallis with Dunn’s test. For **f, g, j** and **k**, comparisons were performed using an Ordinary One-Way ANOVA with Fisher’s LSD. ns = not significant; * = P-value<0.05; ** = P-value<0.01; *** = P-value<0.001; **** = P-value<0.0001. Error bars represent□±□SEM.

Further, assessment of the overall brain structure, specifically the cortex region, showed that both *Smad4-* and *Eng*-iEC-LOH brains exhibited significantly shorter midline lengths compared to all the other genotypes (Fig 7c,d,f,h,j). Measurement of the cortex area indicated no difference between the Eng models; however, *Smad4*-iEC-LOH brains showed a significant decrease in the cortical area when compared to *Smad4*-iECKO and corresponding control brains (Fig 7c,d,g,h,k). Thus, the data reveal that the LOH models can survive for over 2 months and are more severely affected than iECKO counterparts at these induction levels, strongly suggesting that mosaic LOH leads to progressive vascular and structural brain pathology that eventually limits long-term viability.

To better assess the vascular defects in the P70 mice, blue latex dye perfusion experiments were conducted, and vessel diameters and the presence of latex-filled veins and latex leakage was quantified (Fig 8a,b). In the *HHT-*iEC-LOH models, substantially more latex-filled veins were observed compared to all corresponding experimental groups, with the *HHT*-iECKO brains also showing a significant presence of latex labeled veins compared to the control groups (Fig 8c-e,k). Moreover, the average number of dye-filled veins was similar between the Smad4 and Eng LOH models (Fig 8e,k). In terms of generalized leakage (asterisks), *Smad4*-iEC-LOH mice displayed more latex leakage than both controls (Fig 8f), while *Eng-*iEC-LOH mice only had more generalized leakage than *Eng^f/f^* brains (Fig 8l). Interestingly, in both models, the average number of latex-filled veins and latex leakage prevalence were lower than the amounts observed at P7 suggesting that the vascular phenotypes may have improved over time.

**Fig. 8.**
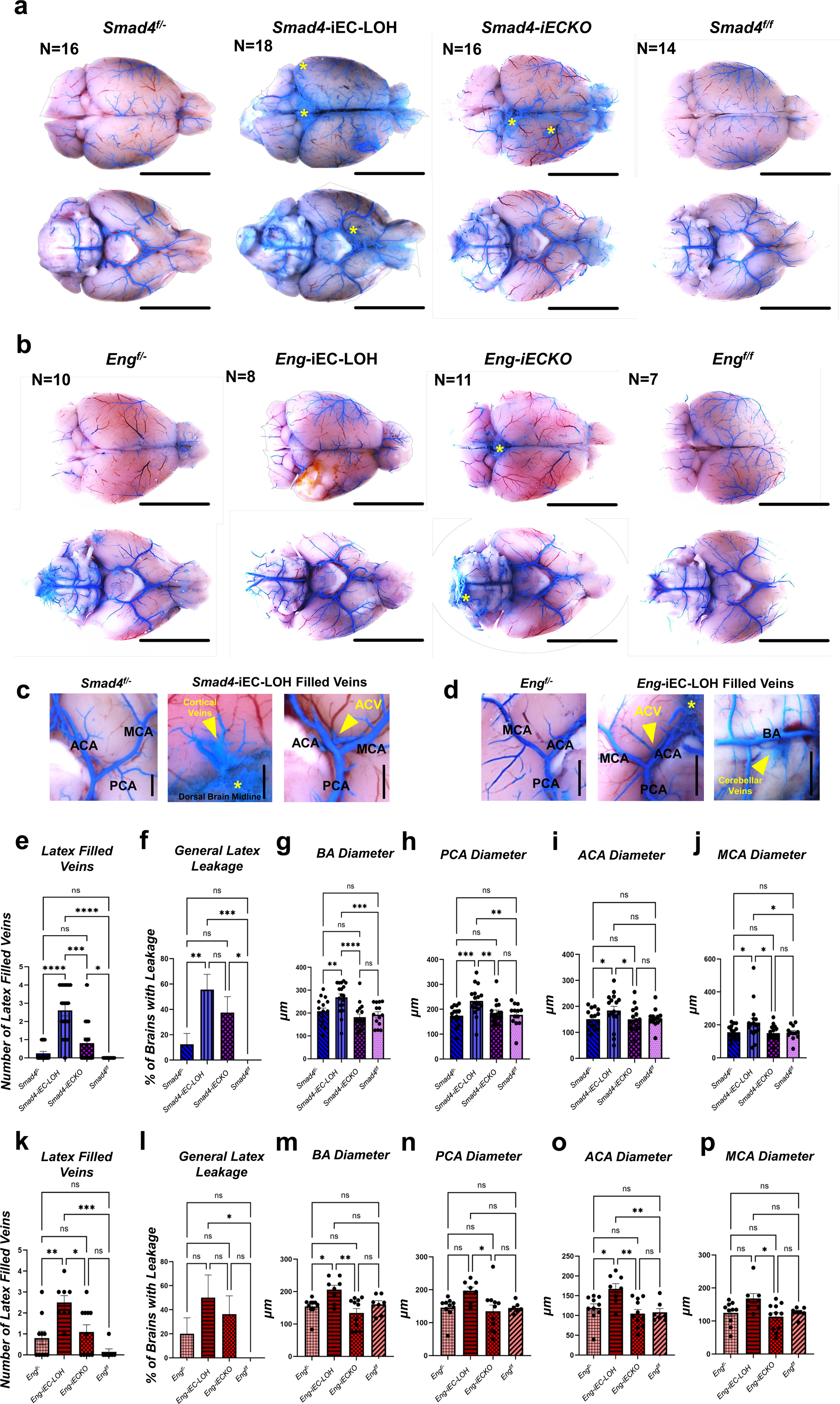
Latex perfusions reveal persistent vascular dysregulation into adulthood. **a, b.** Bright field images of brains from *Smad4 and Eng* models perfused with blue latex. Dorsal (top) and ventral (bottom) views are shown. Scale bars = 5000 µm. **c, d.** Representative close-up images of latex-filled vessels in *Smad4-*iEC-LOH, *Smad4*^f/-^, *Eng-*iEC-LOH and *Eng*^f/-^ brains. Scale bars = 1000 µm. Yellow arrowheads denote veins filled with latex. ACA, anterior communicating artery; MCA, middle communicating artery; PCA, posterior communicating artery; BA, basilar artery; ACV, anterior cerebral vein. **e.** Quantifications show more latex-filled veins in *Smad4-*iEC-LOH brains compared to all other groups. **f.** There is more leakage in *Smad4-*iEC-LOH brains compared to controls, and similar leakage in *Smad4-iECKO* mice. **g-j.** Arterial diameter measurements: **g, h.** BA and PCA diameters are larger in *Smad4-*iEC-LOH brains compared to all other genotypes. **i, j.** ACA and MCA diameters are larger in *Smad4-*iEC-LOH brains compared to *Smad4*-iECKO and *Smad4*^f/-^ mice, while also exhibiting a larger MCA than *Smad4*^f/f^ mice. **k.** Quantifications indicate there are significantly more latex-filled veins in *Eng-*iEC-LOH brains compared to all other groups. **l.** More leakage is observed in *Eng-*iEC-LOH mice compared to *Eng*^f/f^ wild type mice. **m-p.** Arterial diameter measurements: m-p. Diameters were all significantly larger in *Eng-*iEC-LOH brains compared to *Eng*-iECKO mice. **m, o.** Additionally, the BA and ACA dilations were higher in *Eng-*iEC-LOH brains compared to *Eng*^f/-^mice, while also showing an increase in ACA diameter compared to *Eng*^f/f^ brains. For e, f, k, l comparisons were performed with a Kruskal-Wallis test with Dunn’s post-hoc. For g-j and m-p, arterial diameters were measured using ImageJ and compared using an Ordinary One Way ANOVA with Fisher’s LSD. ns = not significant; * = P-value<0.05; ** = P-value<0.01; *** = P-value<0.001; ****P-value<0.0001. Error bars represent□±□SEM.

When analyzing the vessel calibers of multiple arteries, the *Smad4-*iEC-LOH mice, similar to P7, had significantly more dilated basilar and posterior communicating arteries than controls (Fig 8g,h; Sup Fig 5h,i). Furthermore, this increase was also significantly higher than *Smad4-iECKO* mutants at P70 (Fig 8g,h). The anterior communicating artery of *Smad4-*iEC-LOH mice was more dilated than *Smad4^f/-^* and *Smad4-iECKO* mice (Fig 8i), which showed no differences in sizes between all groups at P7 (Sup Fig 5g). Finally, unlike at P7, an increased dilation was observed at P70 in the middle communicating artery (Fig 8j; Sup Fig 5f). In all, the arterial vessel enlargements appeared more pronounced in Smad4 LOH mice at P70 than P7. At P70, *Eng*-iEC-LOH diameter increases were not as prominent as those seen at P7. The posterior and middle communicating artery diameters of *Eng-*iEC-LOH mice were only significantly higher than *Eng-iECKO* mice and comparable to controls (Fig 8n,p; Sup Fig 5n,q). The basilar artery diameter of *Eng*-iEC-LOH mice was increased compared to *Eng^f/-^* and *Eng-iECKO* mice (Fig 8m), whereas in the LOH model at P7 it was significantly larger than all genotypes (Sup Fig 5p). Finally, the anterior communicating artery dilation was significantly higher in *Eng-*iEC-LOH mice compared to all other groups (Fig 8o), which was only larger than the control groups at P7 (Sup Fig 5o). Taken together, it appears that the vessel enlargements are sustained into adulthood, albeit with shifting regional sensitivities, further distinguishing the LOH-driven pathology from the milder phenotypes observed in standard iECKO mice.

## Discussion

Current mouse models of HHT have been integral to progressing the field towards a cure, providing essential insights into the complex disease pathogenesis and the molecular mechanisms underlying vascular malformations. In particular, the homozygous inducible EC-specific knockout (iECKO) mouse models have allowed researchers to investigate the effects of TGFβ signaling loss in a manner relevant to HHT, subsequently becoming the current standard for studying this disease. The *HHT*-iEC-LOH mouse models were designed to complement these existing studies by offering an avenue to examine disease mechanisms within a heterozygous loss of function background. This approach more closely recapitulates the genetic landscape of HHT patients compared to traditional iECKO models. To define the unique utility of these new LOH models, this study sought to comprehensively characterize their vascular phenotypes in comparison to iECKO mouse models and corresponding control genotypes. In both the *Smad4*- and *Eng*-iEC-LOH models, we observed prominent retinal and cerebral vascular defects. Notably, vasculature disruption was more pronounced in the *HHT*-iEC-LOH models than in the *HHT-iECKO* models. While this increased severity was not surprising, given that LOH models initiate from a germline heterozygous null background, it highlights the robustness of this genetic setting in driving the formation of vascular malformations and barrier defects. Therefore, the newly developed LOH models provide a new platform to interrogate the mechanisms of stochastic lesion formation and evaluate therapeutic interventions in a context that more accurately mirrors the genetic landscape and progressive nature of human HHT.

A major benefit of the LOH models is their capacity to better elucidate the mosaic nature of AVM formation. Previous studies utilizing iECKO models indicated that AVMs are mosaic, comprised of both mutant and non-mutant cells[71, 72], though one study suggested that AVMs arise via clonal expansion[73]. However, these homozygous models, in which ECs either harbor HHT genes with complete loss of function or remain wildtype, do not accurately reflect the LOH phenomena observed in humans. Unlike iECKO models, HHT patients carry one inactive germline allele in all cells, while only a subset of ECs experiences the biallelic loss that leads to complete loss of function; the remaining ECs remain heterozygous. Indeed, HHT patient malformations have been shown to consist of ECs with biallelic loss interspersed with germline heterozygous cells[22–26]. Our results with *HHT*-iEC-LOH mouse models demonstrate that, similar to HHT patients, AVMs are comprised of a mosaic mixture of heterozygous and LOH cells. Furthermore, we observed that a threshold of approximately 65-70% of LOH cells was associated with AVM onset in both models. However, as little as 10% and 38% of LOH ECs in Smad4 and Eng LOH retinas, respectively, appeared to be sufficient to drive an AVM. In contrast, the reported level of biallelic loss in telangiectasias and AVMs from HHT patients is significantly lower, at approximately 2-8% of the ECs[22–24]. This numerical gap may reflect the difference between the acute, synchronized induction of biallelic loss via tamoxifen in our murine models and the gradual, sporadic accumulation of somatic mutations over decades in HHT patients, which may permit lesion maturation with a lower initial mutation burden. However, as noted by several groups, technical challenges, such as tissue processing and the inherent difficulty of detecting rare somatic mutations in genomic DNA, have likely impeded the ability to capture the full extent of biallelic loss in patient samples[22, 74]. Thus, the low frequency of biallelic loss currently reported in HHT patient lesions is likely an underestimation. Consequently, it will be critical to characterize the LOH:non-LOH EC composition of vascular malformations as advances in single-cell sequencing and tissue isolation protocols evolve. Based upon our experiments, we would expect that vascular malformations in HHT patients require a critical mass of LOH cells to trigger lesion maturation, even if the initiating proportion of mutant cells is relatively small.

Building on these findings, we used the confetti reporter system to track distinct recombination events and determine how they contribute to AVM development. Based on the diversity of fluorescent markers observed, it appears that *Eng*-iEC-LOH mice require a higher number of distinct mutation events to drive lesion formation, whereas *Smad4*-iEC-LOH mice require fewer instances of biallelic loss. Interestingly, the majority of these recombinant events were localized near the feeding vein, suggesting a venous origin for these lesions – an observation consistent with previous iECKO studies[71, 75]. In both LOH models, the presence of multiple fluorescent markers within a single AVM indicates that lesion development is not strictly the result of a single clonal event. However, it is noteworthy that LOH and non-LOH cells were rarely intermingled; instead, they tended to form discrete clusters. Future research utilizing single-cell and spatial transcriptomics may further shed light on the complex dynamics between LOH and neighboring heterozygous ECs during lesion maturation. Lastly, it will be important to reconcile these findings with the clinical data where typically only a single somatic mutation is detected in lesions from HHT1 (ENG) and HHT2 (ACVRL1) patients[22, 24, 25]. While this clinical observation may support a clonal expansion-based mechanism, at least one case of JP-HHT has revealed two distinct somatic alterations causing biallelic loss of SMAD4 in a single AVM[23]. Advancements in somatic mutation detection will be essential to determine if multiple hits are a general requirement for lesion formation or if the mechanism varies by HHT type. Overall, our data suggest that more than one somatic mutation may be necessary to generate a full-scale AVM, though the required threshold of “hits” may differ between telangiectasias and larger malformations.

Fluorescent-tagged tracers are established tools for evaluating blood brain barrier (BBB) and blood retinal barrier (BRB) defects, which make them ideal for assessing vascular integrity in HHT. For instance, in BMP9/10 antibody blocking and genetic models of HHT, 1 kDa tracer and cadaverine leakage have been observed in the brain and retina of mutant mice, alongside RBC extravasation in the retina[68, 69]. While similar leakage of larger sized tracers has been shown in *in vitro* HHT models[76, 77], there remains a lack of *in vivo* research detailing the extent of vascular permeability alterations across different HHT genotypes – a critical gap in understanding the pathophysiology of vascular barrier dysfunction. Using the *HHT*-iEC-LOH mouse models, this study demonstrated the extravasation of high molecular weighted tracers in the brain, as well as RBC presence outside the retinal vasculature. RBC leakage was observed within hemorrhages, proximal to AVMs, and notably surrounding relatively normal capillary beds. This suggests that loss of barrier function is a widespread phenomenon throughout the microvasculature, even in vessels without overt malformations. These data may provide a mechanistic basis for the chronic bleeding and epistaxis (nosebleeds) that HHT patients experience in the smaller vessels of the nasal and gastrointestinal mucosa. Such persistent leakage likely contributes to negative health outcomes beyond major malformations by dysregulating oxygen and nutrient exchange. Intriguingly, while the overall severity of leakage appeared more pronounced in the *Smad4*-iEC-LOH model, *Eng*-iEC-LOH mice showed a higher frequency of larger tracer escape. It would be of great interest to determine if these findings correlate with the bleeding severity and phenotypes in HHT1 versus JP-HHT patients. Collectively, these results underscore that HHT is fundamentally associated with compromised vessel integrity, suggesting that therapies targeting vascular stability may be as crucial as those targeting the malformations themselves.

A distinct advantage of the *HHT-*iEC-LOH models is their capacity to develop robust, consistent HHT pathology while maintaining long-term viability into adulthood following early induction of LOH. In contrast, iECKO mice treated with equivalent low doses show minimal vascular defects. To drive severe malformations in the iECKO models, significantly higher doses of tamoxifen are required; however, these high-dose regimens do not accurately reflect the stochastic “second hit” nature of human HHT. Furthermore, the doses necessary to produce extensive lesions in iECKO mice often result in early lethality, with mice seldom surviving beyond 10 days of age[30, 40]. Significantly, tamoxifen induction in adult *HHT*-iECKO mice leads to few malformations because they lack the specific developmental, angiogenic time frame present in neonates that is required to trigger lesion formation in a homozygous null context[29, 44]. Consequently, there have been fewer studies investigating the long-term progression of these vascular phenotypes. Most existing research focuses on the prevention of lesion development rather than assessing whether therapeutic interventions can induce AVM regression – a distinction of critical importance to patients who already harbor established lesions. To circumvent these limitations, several groups have utilized secondary insults, such as ear wounding and stereotaxic injection of VEGF or LPS in the adult brain with Cre recombinase, to create a pro-angiogenic environment, or have employed organ- and EC-specific Cre drivers to induce gene deletion at neonate stages and restrict vascular phenotypes to specific [39–42, 46, 78, 79]. However, although useful, these strategies preclude a whole-body assessment of HHT vascular phenotypes, which is essential for understanding the tissue-specific effects in a systemic context. In contrast, our data show that with a low tamoxifen dose, *HHT-*iEC-LOH mice develop significant lesions and hemorrhaging yet survive up to 10 weeks. While further assessment is required to confirm the presence of lesions across various organ systems, these LOH models provide a unique platform for longitudinal studies. This will facilitate research into key timepoints of disease progression and the long-term efficacy of therapeutics in a model that more closely recapitulates the genetic makeup of HHT patients.

By maintaining a heterozygous background while inducing mosaic LOH in ECs, these models provide a platform to observe systemic changes and cellular cross-talk in a way previously not possible. Utilizing Cre reporters such as *mTmG* will allow future studies to investigate the influence of LOH ECs on their non-LOH neighbors and auxiliary cell types. Indeed, it remains unclear how mural cells and the surrounding mesenchyme (which also harbor the germline mutations in this model) respond to focal biallelic loss and/or contribute to HHT lesion formation and maturation; these models offer the precision required to study those dynamics. Furthermore, the ability to mimic the genetic landscape of HHT patients, whereby mutant ECs are embedded within a heterozygous environment, is essential for understanding potential paracrine signaling involved in driving lesion development and maturation. Beyond mechanistic insights, the adult viability of these LOH models enables longitudinal monitoring of disease progression and testing of therapeutics at distinct developmental stages. Such studies will be critical for determining the optimal windows for clinical intervention. Ultimately, this work expands the HHT research repertoire by providing two mouse models that better recapitulate patient biology, marking a significant step forward in the pursuit of targeted treatments and a cure.

## Statements and Declarations

The authors declare no competing interests.

## Acknowledgements

The authors are grateful to Hua Su for transferring the *Eng* conditional floxed mice from UC San Francisco in coordination with Helen Arthur. The authors would also like to thank Angela Crist for her early contributions to the development of the *Smad4-*iEC-LOH mice.

The authors wish to thank the Tulane University Department of Comparative Medicine staff, especially the veterinarians and technicians, for their wonderful support with mouse husbandry.

The authors would like to acknowledge their funding sources. The work shown in this paper has been funded by various grants through the National Institute of Health, National Heart, Blood, and Lung Institute and the Eye, Ear, Nose and Throat Foundation of New Orleans (EENT). These include: F31-HL174077 (APB), R01-HL163196 (SMM), R01-HL139713 (SMM) and EE240902 (SMM).

Figure diagrams were created with Biorender.com.

## Notes

### Competing Interest Statement

The authors have declared no competing interest.

### Summary of Updates

This revision includes Figure Legends for the main figures, which were missing in the original submission.

