## Supplemental files for "Modeling Somatic Second-Hit Mutations in Novel Mouse Models of Hereditary Hemorrhagic Telangiectasia"

### Supplemental Figures

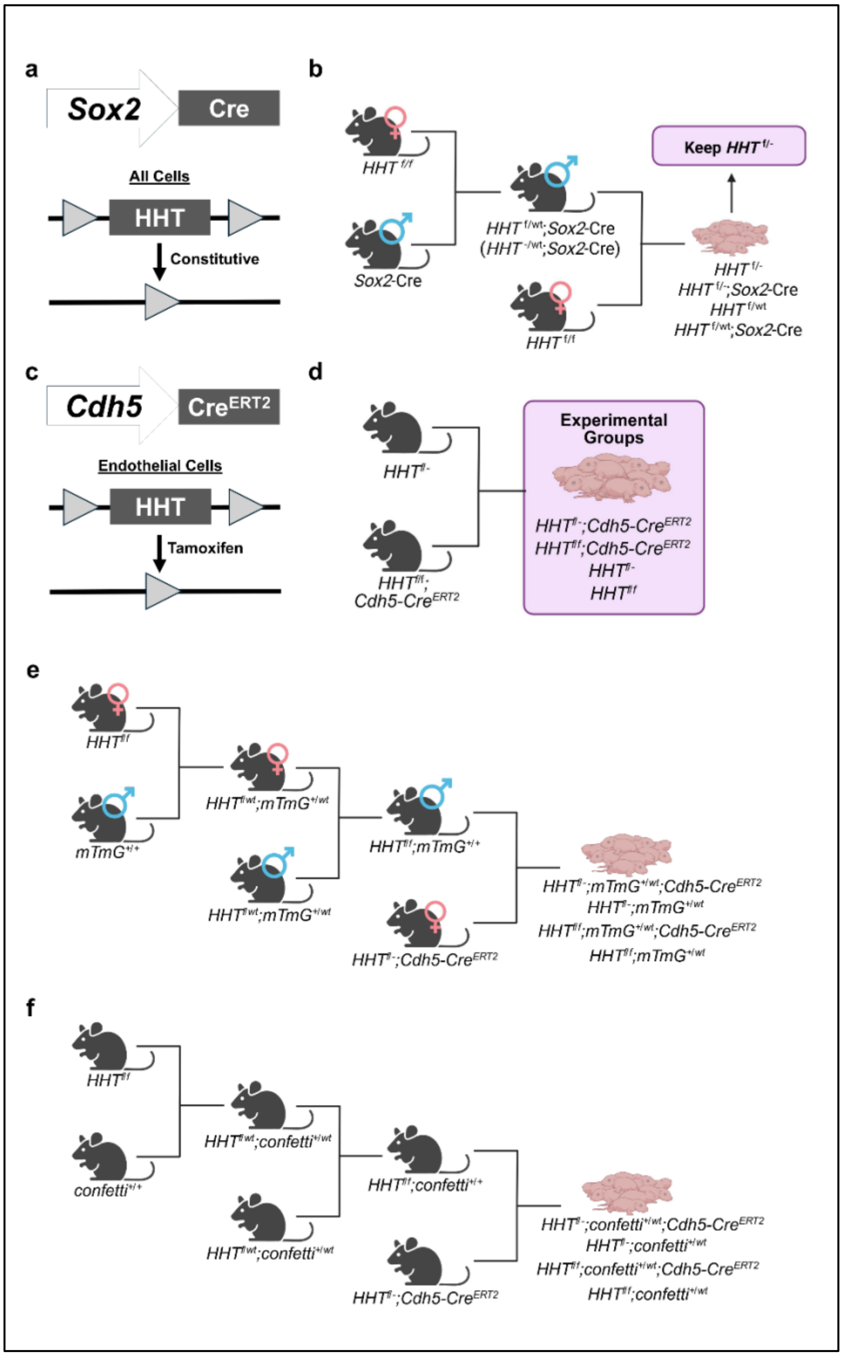


**Supplementary Fig. 1 Mating strategies to generate variations of *HHT*-iEC-LOH mouse models using the Cre-LoxP system**

**a.** The *Sox2*-Cre construct is endogenously active in germline cells, resulting in the excision of the floxed *HHT* allele in all cells of progeny with Cre. **b.** Breeding scheme to generate *HHT*$^{f/-}$ mice to be used in experimental matings. *HHT*$^{-/wt}$;*Sox2*-Cre males were mated to *HHT*$^{f/f}$ females to produce *HHT*$^{f/-}$ mice lacking *Sox2*-Cre. Only *Sox2*-Cre male mice were used to avoid maternal germline inheritance issues. **c.** The *Cdh5*-Cre$^{ERT2}$ construct is tamoxifen-inducible, specifically in ECs. **d.** Experimental mating strategy for LOH mice. Breeding scheme produces four possible genotypes: *HHT*-iEC-LOH (*HHT*$^{f/-}$;*Cdh5*-Cre$^{ERT2}$), *HHT*-iECKO (*HHT*$^{f/f}$;*Cdh5*-Cre$^{ERT2}$), *HHT*$^{f/-}$ and *HHT*$^{f/f}$. **e.** Breeding scheme to generate *HHT*$^{f/f}$;*mTmG*$^{+/+}$ mice, which were used to mate with *HHT*-iEC-LOH mice to generate the four possible genotypes with *mTmG*. Since *mTmG*$^{+/+}$ females did not adequately care for pups, males of this genotype were utilized. **f.** Breeding scheme to generate *HHT*$^{f/f}$;*confetti*$^{+/+}$ mice for experimental mating with *HHT*-iEC-LOH mice to generate the four possible genotypes with *confetti*.

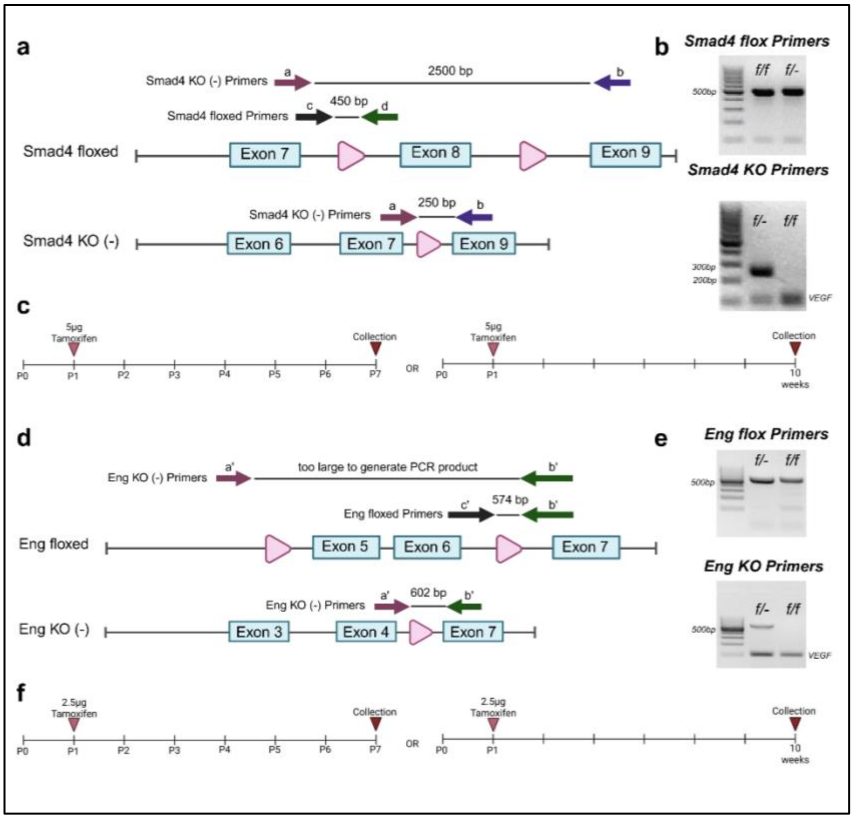


**Supplementary Fig. 2 Genetic transgenic constructs and genotyping strategies for the *HHT* models**

**a.** Genetic construct of the *Smad4* floxed allele. LoxP sites flank Exon 8, which is excised upon Cre-mediated recombination using tamoxifen. Approximate binding sites of PCR primers and expected product sizes are displayed. **b.** Representative PCR genotyping results. *Smad4* floxed primers detect the presence of a loxP site (intact floxed Exon 8) and yield a 450 base pair (bp) band in both flox/flox or flox/- backgrounds. *Smad4* knockout (-) primers identify the excision of Exon 8, resulting in one 250 bp band when the null knockout (-) allele is present; these primers also produce an approximately 2500 bp band in flox/flox mice. Gels were resolved alongside a 100 bp ladder. **c.** Tamoxifen treatment strategy for *Smad4*-iEC-LOH mice. A dose of 5 µg of tamoxifen was fed to mice at P1, and tissues were collected at P7 or P70 (10 weeks). **d.** Genetic construct of the *Eng* floxed allele. LoxP sites flank Exons 5 and 6. Approximate binding sites of PCR primers and product sizes are displayed. **e.** Representative PCR genotyping results. *Eng* floxed primers detect a single loxP site (intact floxed Exons 5 and 6) and yield a 574 base pair (bp) band in both flox/flox or flox/- backgrounds. *Eng* knockout (-) primers identify the absence of Exons 5 and 6 and yield a 602 bp band only when the null knockout (-) allele is present. The *Eng* knockout primers do not produce a band in the presence of the floxed allele. Gels were resolved alongside a 100 bp ladder. **f.** Tamoxifen treatment strategy for *Eng*-iEC-LOH mice. A does of 2.5 µg of tamoxifen was fed to mice at P1, and tissues were collected at P7 or P70 (10 weeks).

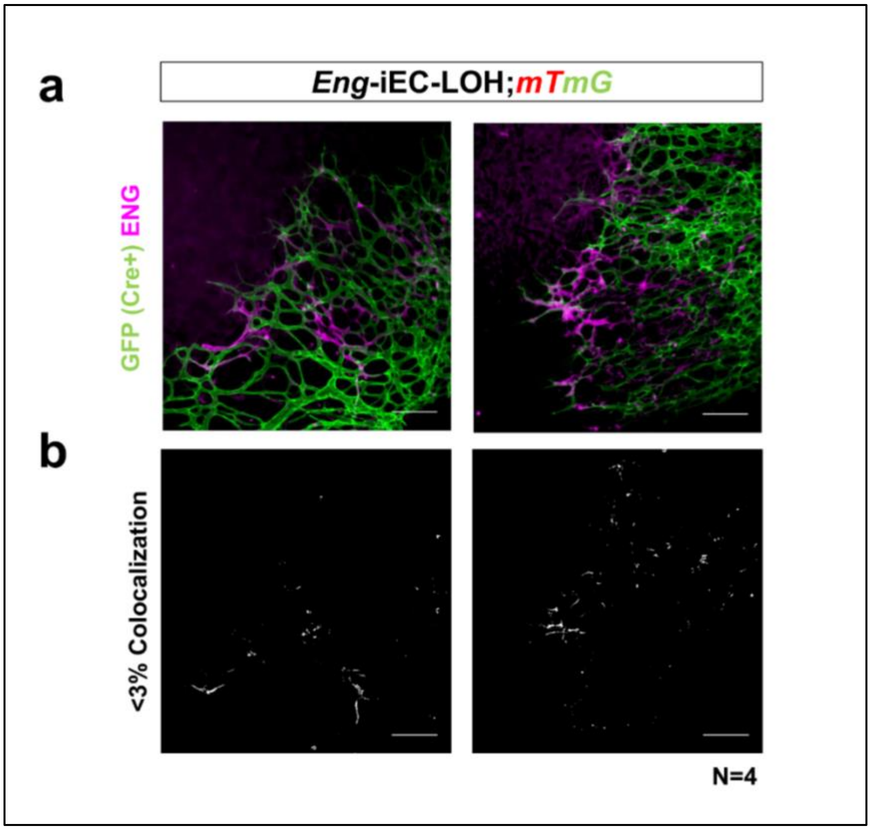


**Supplementary Fig. 3 Validation of ENG knockout in mTmG retinas**

**a.** Immunostaining for ENDOGLIN (ENG/CD105) in *Eng*-iEC-LOH;*mTmG* P7 retinas confirms gene deletion in eGFP-positive ECs. Consistent with successful knockout, ENG-positive ECs lack eGFP expression, whereas eGFP-expressing cells show a loss of ENG signal. Scale bars = 100 µm. **b.** Quantitative colocalization analysis using ImageJ demonstrates that less than 3% of the total fluorescent area shows overlap between ENG and eGFP signals. These data validate eGFP as a reliable surrogate marker for biallelic *Eng* deletion in the *Eng*-iEC-LOH model.

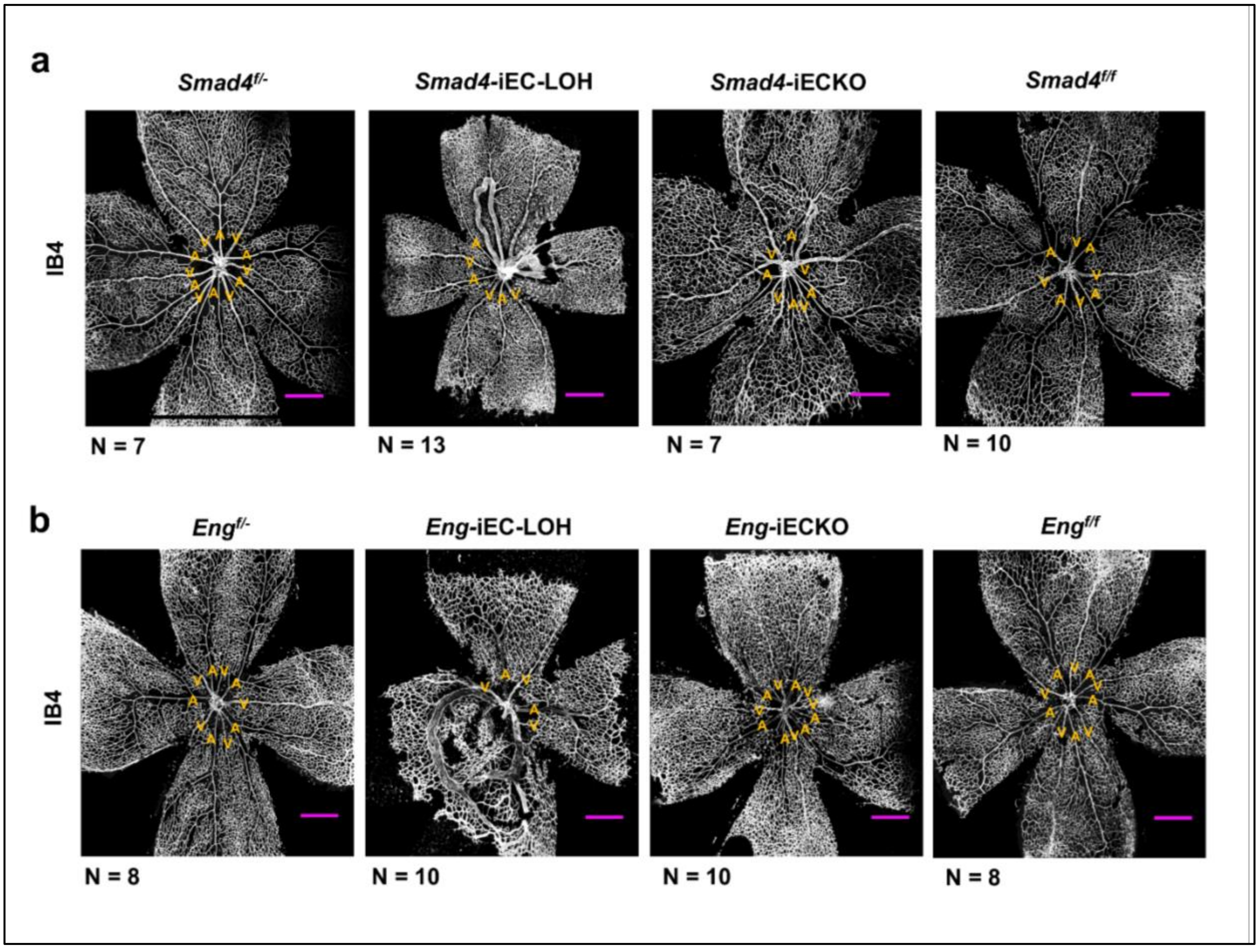


**Supplementary Fig. 4 Global retinal vasculature defects in *HHT*-iEC-LOH mice**

**a, b.** Representative whole-mount images of P7 retinas from all *Smad4* and *Eng* models portrayed in Figure 3; immunostained with ISOLECTIN B4 (IB4) to visualize the vasculature. These low-magnification overviews highlight the systemic distribution of vascular malformations and remodeling defects across the different genotypes. Scale bars = 500 µm. A, artery; V, vein.

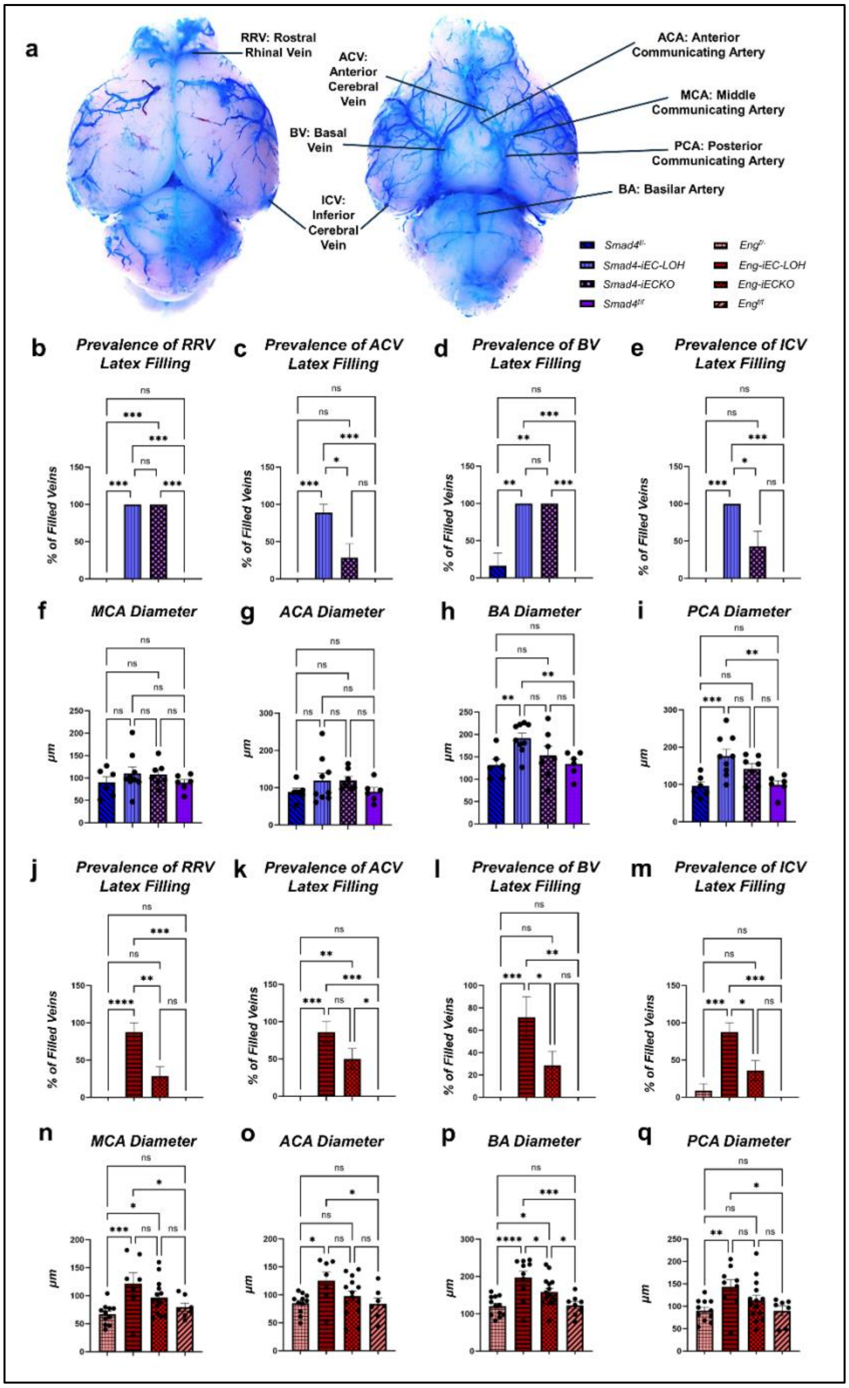


**Supplementary Fig. 5 Diameter changes can be seen in *HHT*-iEC-LOH brains at P7 a.** Diagram of the four major veins characterized for blue latex presence and four major arteries which diameters were measured. **b.** The prevalence of RRV filling was higher in both mutant mice compared to controls. **c.** The prevalence of ACV filling was higher in *Smad4*-iEC-LOH mice compared to all others. **d.** The prevalence of BV filling was higher in both mutant mice compared to controls. **e.** The prevalence of ICV filling was higher in *Smad4*-iEC-LOH mice compared to all others. **f.** The MCA diameters were similar in all four *Smad4* models. **g.** The ACA diameters were similar in all four *Smad4* models. **h.** The BA diameter of *Smad4*-iEC-LOH mice is increased compared to controls. **i.** The PCA diameter of *Smad4*-iEC-LOH mice is increased compared to controls. **j.** The prevalence of RRV latex filling was highest in *Eng*-iEC-LOH mice. **k.** The prevalence of ACV filling was higher in *Eng*-iEC-LOH mice compared to controls. **l.** The prevalence of BV latex filling was highest in *Eng*-iEC-LOH mice. **m.** The prevalence of ICV latex filling was highest in *Eng*-iEC-LOH mice. **n.** The MCA diameters were higher in *Eng*-iEC-LOH models. **o.** The ACA diameters were higher in *Eng*-iEC-LOH mice compared to controls. **p.** The BA diameter of *Eng*-iEC-LOH mice is increased compared to all mice. **q.** The PCA diameter was higher in *Eng*-iEC-LOH mice compared to controls. For b-e and j-m, comparisons were performed using a Kruskal-Wallis test with Dunn's post-hoc. For f-i and n-q, diameters were compared using a Ordinary One-Way ANOVA with Fisher's LSD. ns = not significant; * = P-value<0.05; ** = P-value<0.01; *** = P-value<0.001; **** = P-value<0.0001. Error bars represent ± SEM.

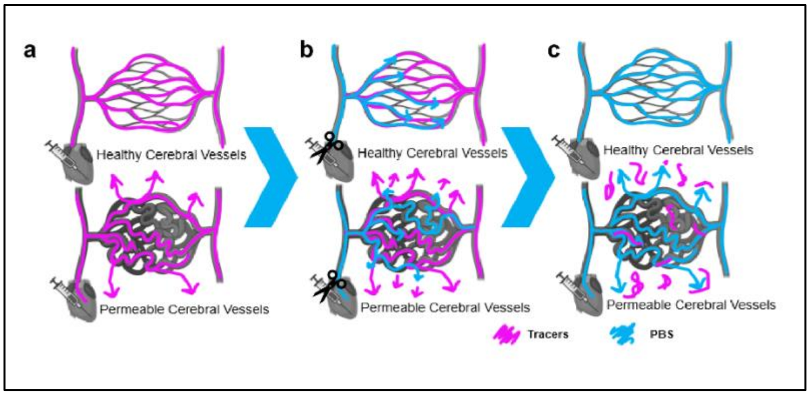


**Supplementary Fig. 6 Schematic of the Tracer Perfusion Workflow**

**a.** Tracers (pink) are transcardially perfused into the systemic circulation. In healthy cerebral vessels, tracers are confined to the vascular lumen. In permeable vessels, tracers extravasate into the surrounding parenchyma. **b.** Following tracer circulation, the right atrium of the heart is clipped, and PBS is perfused to flush the vasculature. In healthy vessels, PBS clears tracers from the vessels. In permeable vessels, tracers that leaked into the interstitial spaces remains sequestered outside the vasculature. **c.** After complete PBS perfusion, healthy cerebral vessels appear completely cleared of tracers, with no tracers in the parenchyma. Conversely, permeable and disorganized vessels exhibit residual tracer signal in the extravascular space.

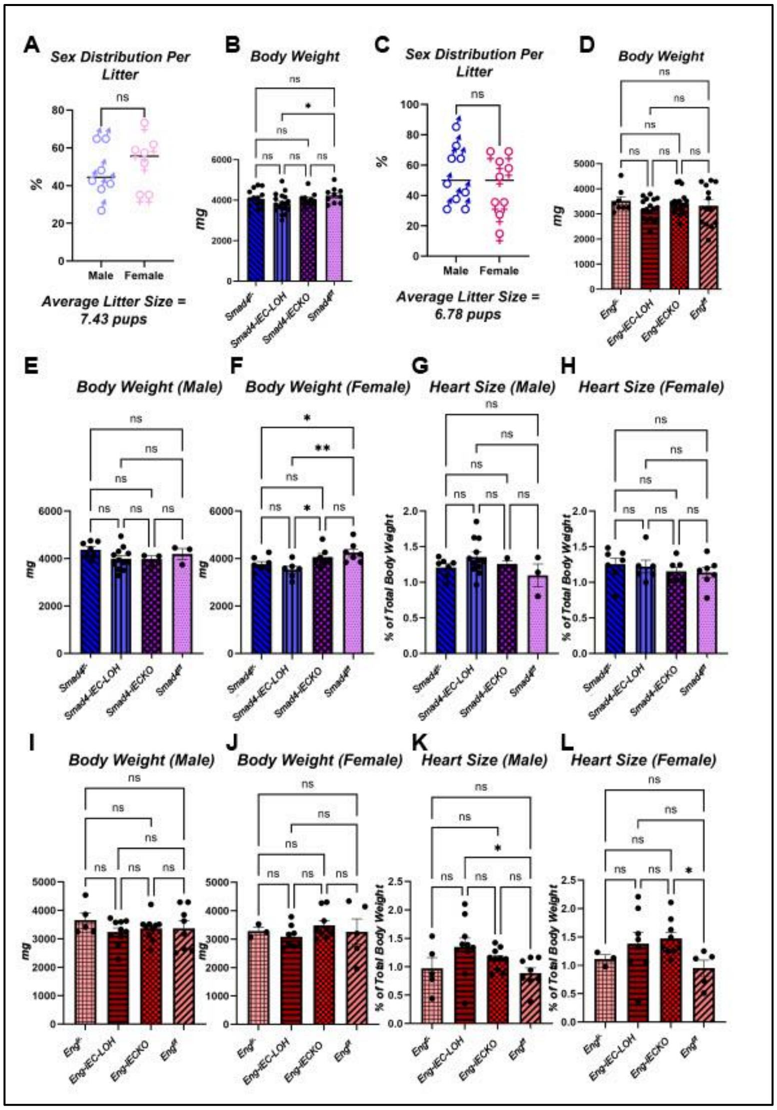


**Supplementary Fig. 7 Body weight and cardiac morphometrics in *HHT*-iEC-LOH mouse models**

**a,c**. Sex distribution ratios in *Smad4* (a) and *Eng* (c) litters exhibit an equal distribution between males and females (unpaired t-test. ns = not significant). **b, d.** Analysis of P7 body weights (sexes combined) showed a reduction in *Smad4*-iEC-LOH pups compared to *Smad4*$^{f/f}$ controls (b), whereas no significant weight differences were observed in the *Eng* cohorts (d). **e-h.** Analysis of the *Smad4* model. **e, f.** Sex-disaggregated body weight analysis reveals no significant differences among *Smad4* males (e). However, *Smad4*-iEC-LOH females weigh significantly less than both *Smad4*$^{f/f}$ and *Smad4*-iECKO mice (f). *Smad4*$^{f/-}$ females also show a significant weight reduction compared to *Smad4*$^{f/f}$ controls. **g, h.** No significant differences in heart weight (normalized as a percentage to total body weight) are observed in either male (g) or female (h) *Smad4* mice across genotypes. **i-l.** Analysis of the *Eng* model. **i, j.** No significant differences in body weight are observed among *Eng* males (i) or females (j). **k, l.** Cardiac morphometrics reveal that *Eng*-iEC-LOH male hearts are significantly enlarged compared to *Eng*$^{f/f}$ controls (k), while *Eng*-iECKO female hearts are significantly larger than *Eng*$^{f/f}$ controls (l). For panels b, d, and e–l, data were analyzed using Ordinary One-Way ANOVA with Fisher's LSD post-hoc test. ns = not significant; $^*P < 0.05$; $^{**}P < 0.01$.
